# Hippocampal neuroinflammation causes sex-specific disruptions in action selection, food approach memories, and neuronal activation

**DOI:** 10.1101/2024.05.19.594460

**Authors:** Kiruthika Ganesan, Sahar Ghorbanpour, William Kendall, Sarah Thomas Broome, Joanne M. Gladding, Amolika Dhungana, Arvie Rodriguez Abiero, Maedeh Mahmoudi, Alessandro Castorina, Michael D. Kendig, Serena Becchi, Veronika Valova, Louise Cole, Laura A. Bradfield

**Affiliations:** School of Life Sciences, Faculty of Science, University of Technology Sydney, Sydney, New South Wales 2007, Australia; Centre for Neuroscience and Regenerative Medicine, St. Vincent’s Centre for Applied Medical Research, St. Vincent’s Health Network, Sydney, New South Wales 2010, Australia; School of Psychology, Faculty of Science, University of Sydney, New South Wales 2006, Australia; Institute of Cell and Tissue Culture Technologies, Department of Biotechnology, BOKU University, Vienna, Austria; Sorbonne Université, Institut du Cerveau - Paris Brain Institute - ICM, Inserm, CNRS, APHP, Hôpital de la Pitié Salpêtrière, 75013 Paris, France; Decision Neuroscience Laboratory, School of Psychology, University of New South Wales Sydney, Sydney, New South Wales 2052, Australia; Teva Pharmaceuticals, Sydney, New South Wales 2113, Australia; School of Medical Sciences, Faculty of Medicine and Health, University of Sydney, New South Wales 2050, Australia

**Author notes:** **Corresponding author: Dr. Laura Bradfield. Email:** X: Bradfield_Neuro School of Life Sciences, Faculty of Science, University of Technology Sydney, NSW 2007.

**Keywords:** Hippocampus, sex differences, neuroinflammation, microglia, astrocytes, instrumental action selection, Pavlovian memory, goal-directed action, locomotor activity

## Abstract

Hippocampal neuroinflammation is present in multiple diseases and disorders that impact motivated behaviour in a sex-specific manner, but whether neuroinflammation alone is sufficient to disrupt such behaviour is unknown. We investigated this question here using mice. First, the application of an endotoxin to primary cultures containing only hippocampal neurons did not affect their activation. However, when the same endotoxin was applied to mixed neuronal/glial cultures it did increase neuronal activation, providing initial indications of how it might be able to effect behavioural change. We next demonstrated neuroinflammatory effects on behaviour directly, demonstrating that intra-hippocampal administration of the same endotoxin increased locomotor activity and accelerated goal-directed learning in both male and female mice. In contrast, hippocampal neuroinflammation caused sex-specific disruptions to the acquisition of instrumental actions and to Pavlovian food-approach memories. Finally, we showed that hippocampal neuroinflammation had a sexually dimorphic effect on neuronal activation: increasing it in females and decreasing it in males.

## Introduction

Hippocampal neuroinflammation is a common pathological event in diseases such as Alzheimer’s disease^1^, multiple sclerosis^2^, and depression^3^, each of which feature disruptions of motivated behaviour^4–6^, and each of which is more prevalent in females than males^7–9^. It has therefore been speculated that hippocampal neuroinflammation could be the cause of gender-specific alterations in behaviour and cognition observed in these diseases^7,10,11^. However, because each of these diseases presents with additional neuropathological and structural changes/abnormalities, such as the deposition of amyloid plaques and tau tangles in Alzheimer’s^12,13^, demyelination in multiple sclerosis^14^, and frontal lobe atrophy in depression^15^, it is unclear whether neuroinflammation alone is sufficient to cause the behavioural alterations. Here, we tackled this question in a causal manner for the first time, using mice.

There are many similarities in the types of behaviour affected by the aforementioned diseases suggesting that, as the common neuropathological feature, hippocampal neuroinflammation could be the underlying cause. For instance, Alzheimer’s, depression, and multiple sclerosis all present with alterations in food-seeking^16–18^, apathy^10,11,19^, and the ability to carry out daily activities^10,20,21^. Notably, the nature of these effects (e.g. whether the behaviour decreases or increases in propensity) varies widely in a manner that is often associated with gender and/or sex^10,11,16,20,21^ (note that we here use ‘gender’ in reference to the social construct of gender as it applies to humans, and sex in reference to the biological construct as it applies to animals including humans).

Sex also influences neuroinflammation and its impact on the brain. Neuroinflammation is a complex process involving the phenotypic transformation of microglia and astrocytes to a polarised state, which then secrete cytokines^22^ and modify their morphology and function^23,24^, ultimately affecting the activity of neighbouring neurons^25–27^. Given that sex influences the baseline number of glia present in the hippocampus^28,29^, the functional state of those glia^30,31^, and the overall immune response of an organism^32,33^, it is unsurprising to think that neuroinflammation might also lead to sex-specific behavioural effects.

To answer these questions, we first conducted an *in vitro* study. This showed that lipopolysaccharide (LPS) – an endotoxin and neuroinflammatory mimetic – does not alter neuronal activation (c-Fos expression) when applied to hippocampal neuronal monocultures but significantly increases it when applied to neuronal/glial cocultures, particularly in the presence of astrocytes. We subsequently investigated the *in vivo* consequences of hippocampal neuroinflammation by directly injecting LPS into the dorsal hippocampus of female and male mice. We evaluated various instrumental and Pavlovian behaviours, including the propensity to press levers for food, to exert goal-directed control over food-seeking behaviour, and to approach a magazine port associated with food. We further tested the effects on general locomotor activity and anxiety-like behaviours. Our findings revealed a spectrum of behavioural effects produced by hippocampal neuroinflammation, some of which were consistent across sexes and others that were sex-specific. Immunohistochemical analyses produced evidence consistent with neuroinflammation in the hippocampus of both female and male mice (i.e. increased expression of pro-inflammatory cytokine, tumour necrosis factor alpha (TNF-α), as well as altered intensity and morphology of cells positive for ionised calcium binding adaptor 1 molecule (IBA1), a microglial marker, or glial fibrillary acidic protein (GFAP), an astrocytic marker), whereas it produced sexually dimorphic effects on neuronal activation (i.e. c-Fos colocalised with neuronal marker NeuN).

## Results

### Neuroinflammation causes neuronal activation in primary cell cultures, but only in the presence of glia (astrocytes)

Prior to testing whether hippocampal neuroinflammation can alter behaviour, we first sought to identify the potential means by which it could do so. LPS administration is known to cause the release of a range of inflammatory mediators capable of triggering the phenotypic shift of microglia and astrocytes towards pro-inflammatory states, which enables microglial engulfment of foreign pathogen-associated molecular patterns and the activation of reparative processes by astrocytes to restore homeostasis. For this to ultimately alter behaviour, however, it must modulate neuronal activity. This is because glial processes are short and typically confined to their immediate surroundings^34,35^, whereas the neural circuitry underlying motivated behaviours is relatively extensive throughout the brain^36,37^. As a result, hippocampal glia lack the ability to make contact with much of this circuitry, such that any behavioural changes eventuating from altered hippocampal glial function can only occur through their modulation of neurons. Therefore, the first aim of the current study was to determine whether neuroinflammation altered neuronal activation and, if so, whether it relied on modulation by microglia or astrocytes individually to do so, or both together.

To achieve the cell-type precision necessary to answer this question, we used *in vitro* cell culture. As shown in Figure 1A, we applied LPS to primary cell cultures consisting of only hippocampal neurons, to cocultures consisting of neurons and microglia or neurons and astrocytes, and to tricultures consisting of all three cell types. Co- and tricultures were established using monocultured cells at proportions of 2 neurons to 5 astrocytes and/or 1 microglia, and experimental settings were repeated with cells from three biological replicates, each replicated twice (i.e. two technical replicates). Cultures were treated with 1 μg/mL LPS for 24 hours. This concentration was chosen because it has previously been demonstrated to produce a range of neuroinflammatory responses in cell culture^38,39^.

**Figure 1:**
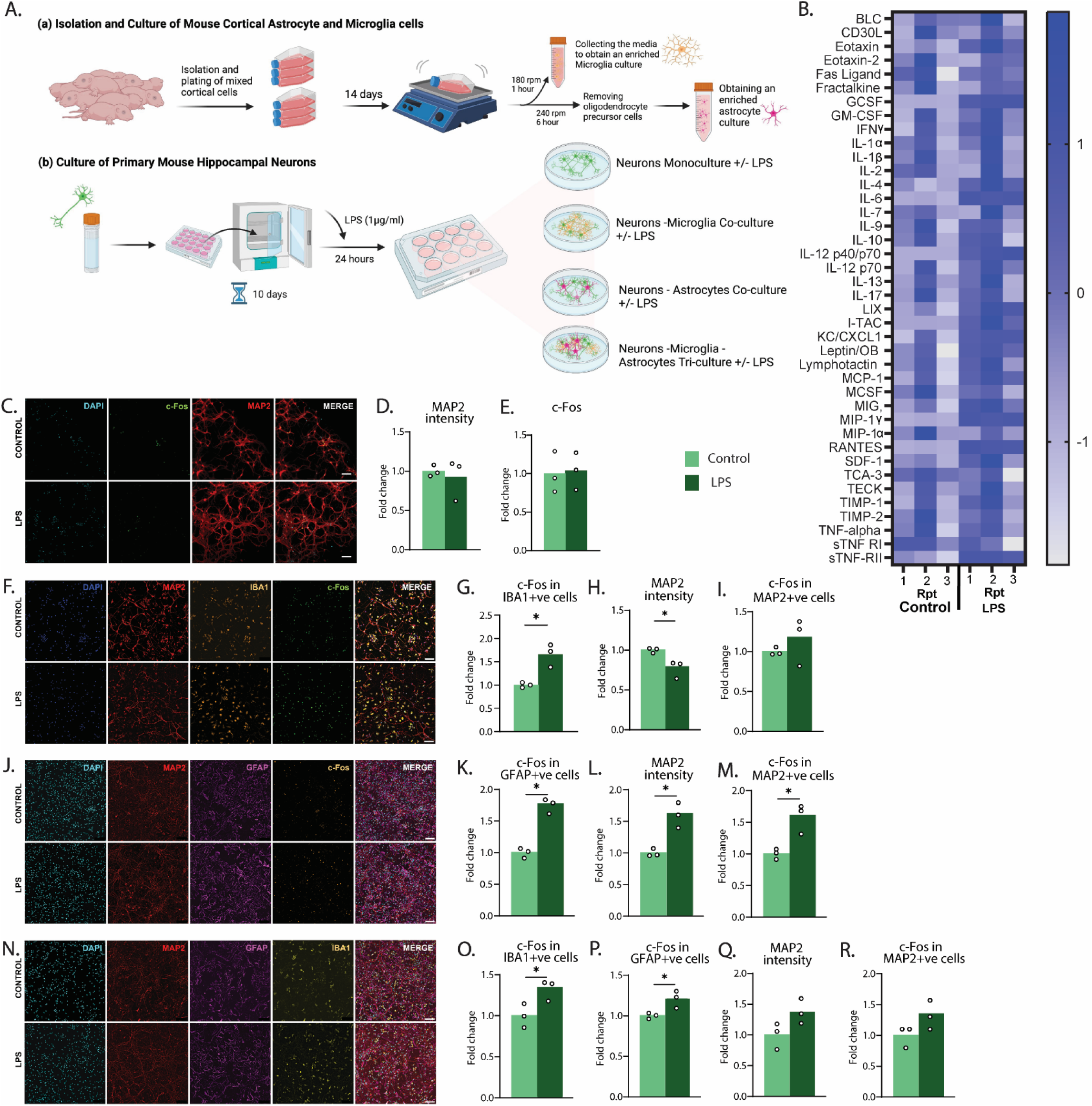
lipopolysaccharide application to primary cultures causes neuronal activation in the presence of astrocytes: (A) a) Schematic representation of the procedure for the collection and processing of mouse hippocampal microglia and astrocytes, and b) for mouse hippocampal neurons, for primary cell culture studies. Created with Biorender.com. (B) heatmap depicting fold change in protein expression of different cytokines from the untreated (control) and LPS-treated neuron, microglia, and astrocyte tricultures. (C,F,J,N) High resolution images showing saline and LPS treated cultures stained with DAPI-nucleus, MAP2-neurons, c-Fos-activated neurons, IBA1-microglia, GFAP-astrocytes (scale bar 100µm) for: (C) neuronal monoculture; (F) neuron microglial coculture; (J) neuron astroglial coculture, (N) neuron microglia and astrocytes triculture. (D,E) Fold change of MAP2 and c-Fos intensity in control (light green) and LPS (dark green) treated neuronal monocultures. (G,H,I) Fold change of MAP2 and c-Fos intensity of neurons and c-Fos intensity of microglial cells in control (light green) and LPS (dark green) treated neuron microglial coculture. (K,L,M) Fold change of MAP2 and c-Fos intensity of neurons and c-Fos intensity of astrocyte cells in control (light green) and LPS (dark green) treated neuron astroglial coculture. (O,P,Q,R) Fold change of MAP2 and c-Fos intensity of neurons and c-Fos intensity of microglial and astrocytic cells in control (light green) and LPS (dark green) treated neuron microglia and astrocytes triculture. For each culture, n = 3 biological replicates (each technically replicated once). *p<0.05.

Untreated cultures were used as controls. We then quantified levels of microtubule-associated protein 2 (MAP2) to identify neurons and neuronal health, as well as expression levels of immediate early gene and neuronal activation marker c-Fos that overlapped with MAP2. For cultures containing microglia we also quantified IBA1, and for cultures containing astrocytes we quantified GFAP, both individually and in conjunction with c-Fos.

Unlike neurons, for which changes in c-Fos expression are typically associated with functional activation^40,41^, the presence of c-Fos in glia can reflect multiple processes including growth, inflammation, and proliferation, among others^41^. Therefore, we took any changes in c-Fos expression that overlapped with MAP2 to indicate functional activation of neurons, whereas changes in c-Fos that overlapped with IBA1 or GFAP were taken to indicate changes in glial activity more generally (although we note that these changes may or may not be related to neuroinflammatory processes).

The purity of microglia and astrocytic cultures is shown in Supplemental Figure 1. Microglial monocultures had at least 87% purity, whereas astrocytic monocultures were minimally 74% pure. We also performed a cytokine assay from the cell supernatant of the tricultures, for which the results are shown in Figure 1B and Supplemental Figure 2. This was done to determine whether there were increases in pro-inflammatory cytokines and determine whether LPS was causing a response that was consistent with inflammation. This was largely the case, as we found that there were increases in the expected cytokines such as IL-6 (Supp Fig. 2F) and IL-12 (Supp Fig. 1G).

For the main analyses, we reported intensity measurements (i.e. mean grey value minus background) for MAP2 alone, or for c-Fos co-localised with MAP2, IBA1, or GFAP. However, graphs that show the quantification of cell counts as well as intensity measurements of IBA1 and GFAP from these same cultures can be found in Supplemental Figure 3.

Figure 1B shows images from a control (top row) and LPS-treated (bottom row) neuronal monoculture. Overall, LPS administration had no effect on any measure when applied to culture containing hippocampal neurons alone. Figures 1D-E show the fold change in MAP2 and c-Fos intensity, respectively, neither of which was affected by LPS treatment, for MAP2, t(4) = .046, p = 0.669, and for c-Fos, t(4) = 0.175, p = 0.87. Because MAP2 is a marker for the entire somatodendritic compartment of a neuron^42^ – a measurement that is captured by our quantification of MAP2 intensity – these results suggest that, not only did LPS have no impact on the number of neurons (as confirmed in Fig. SP3A), but it also did not alter dendritic branching. Nor did LPS alter neuronal activation in this culture, as measured by c-Fos.

Figure 1F depicts images captured from a control (top row) and LPS-treated (bottom row) neuron/microglia coculture. LPS treatment increased microglial activity in this coculture, as expected, because LPS treatment increased c-Fos expression in IBA1-positive cells relative to controls, t(4) = 4.842, p = 0.029 (Fig. 1G). MAP2 expression decreased in LPS-treated cultures relative to controls, t(4) = 3.486, p = 0.0252 (Fig. 1H), but this difference was not attributable to neuronal death as there was no difference in the number of MAP2+ve cells between controls and LPS cultures, t(4) = .561, p = .605 (Fig. SP3D). Rather, this change likely represents dendritic loss. Finally, for this coculture, LPS treatment did not alter neuronal activity because c-Fos expression in MAP2-labelled cells did not differ between groups, t(4) = .999, p = .375 (Fig. 1I).

Figure 1J shows images from a control (top row) and LPS-treated (bottom row) neuron/astrocyte coculture. There was an increase in astrocyte activity caused by the application of LPS because there was an increase in c-Fos in GFAP-positive cells in treated cultures relative to controls, t(4) = 9.229, p = 0.008 (Fig. 1K). This time, however, LPS treatment also increased MAP2 expression relative to controls, t(4) = 4.974, p = 0.0076 (Fig. 1L), and this again appears to be due to a change in dendritic branching rather than the number of neurons, because cell counts for MAP2 were unchanged, t(4) = .326, p = .236 (Fig. SP3G). Importantly, there was an increase in c-Fos in MAP2-labelled cells in the LPS-treated culture, t(4) = 4.04, p = 0.0156 (Fig. 1M), suggesting that there was neuronal activation when LPS was applied to neurons in the presence of astrocytes.

Finally, Figure 1N shows images from a control (top row) and LPS-treated (bottom row) neuron/microglia/astrocyte triculture. In line with the coculture results, both microglial and astrocytic activity increased following treatment with LPS, as there was an increase in c-Fos in both IBA1-positive cells, t(4) = 2.91, p = 0.043 (Fig. 1O), and GFAP-positive cells, t(4) = 3.23, p = 0.0321 (Fig. 1P). This time, however, there was no difference between LPS-treated cultures and controls in overall MAP2 expression, t(4) = 2.208, p = 0.092 (Fig 1Q), and no difference in c-Fos expression in MAP2-positive cells, t(4) = 1.908, p = 0.129 (Fig 1R), despite the directional difference matching that from the astrocyte/neuron coculture (LPS > Control).

Together, these results suggest that LPS-induced neuroinflammation can lead to the activation of hippocampal neurons but only when glia are present, particularly astrocytes.

### Hippocampal neuroinflammation increases the performance of instrumental food-seeking actions in female mice, whereas it accelerates the acquisition of goal-directed action control in mice of both sexes

Following the identification of glial (astrocyte)-modulated neuronal activation as the potential means by which neuroinflammation could cause behavioural change, we next tested how hippocampal neuroinflammation altered motivated behaviour in male and female mice directly. Because the ability to carry out daily activities, and food-seeking in particular, is impaired in many diseases that feature hippocampal neuroinflammation^10,20,21^, we first investigated whether such neuroinflammation could be the cause of disruptions to instrumental (food-seeking) actions or the ability exert goal-directed control over such actions. A timeline for this Experiment is shown in Figure 2A. Mice were aged 9-10 weeks at the beginning of the experiment. Male and female mice each received bilateral injections of either saline (group Sham) or LPS (4µg/µl, group LPS) into dorsal hippocampus (CA1/Dentate gyrus region) prior to the start of behavioural training which consisted of instrumental acquisition, followed by outcome devaluation tests to determine whether the acquired lever press actions were under goal-directed control. This dose (4µg/µl) was chosen because it has previously been shown to produce an increase in gliosis that persisted up to 28 days following intra-hippocampal administration, approximately matching the time-frame required to complete all behavioural procedures in the current study.

**Figure 2:**
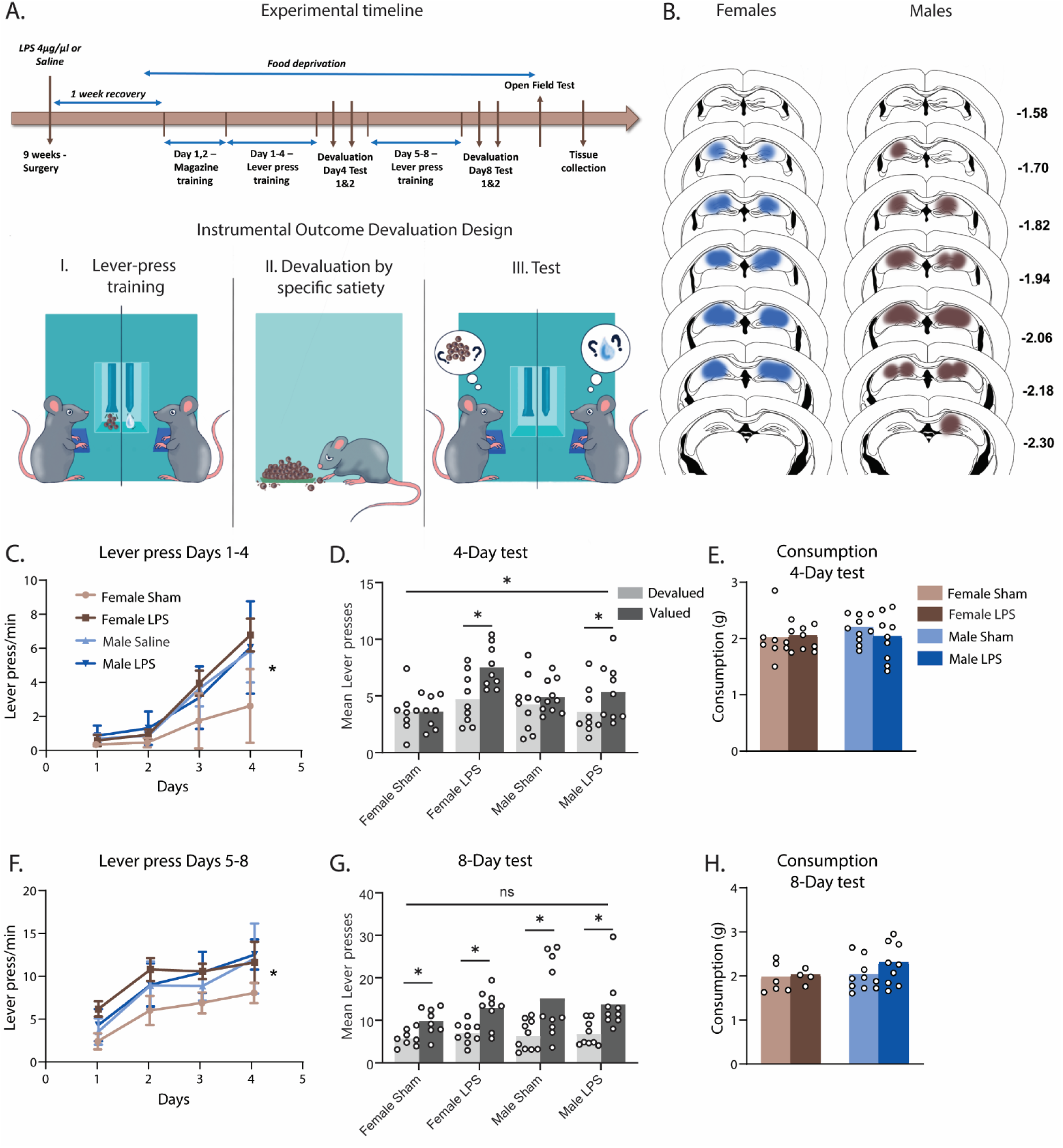
Hippocampal neuroinflammation causes sex-specific effects on instrumental responding but accelerates goal-directed learning in a consistent manner across sexes. (A) Top: Experimental timeline, Bottom: Pictorial representation of the outcome devaluation procedure: 1. Mice are trained to press a left and right lever for pellet and sucrose outcomes (counterbalanced), 2. Mice are prefed to satiety on one of the two outcomes to devalue it, 3. Mice are given a choice between the two levers, but no outcomes are delivered. (B) Representation of LPS placements and spread for Saline and LPS bilateral injections to dorsal hippocampus. (C) Mean lever presses per minute (±SEM) across days 1-4 of lever press training, (D) Mean sum of lever presses during the Day-4 devaluation test, (E) Mean g of reward consumption during prefeeding for Day-4 test, (F) Mean lever presses per minute (±SEM) across days 5-8 of lever press training, (G) Mean sum of lever presses during the Day-8 devaluation test, (H) Mean g of outcome consumption during prefeeding for Day-8 test. *p<0.05.

The design for this experiment is shown in Figure 2A(I-III). Mice were first trained to press two levers for two unique outcomes. Specifically, half of the mice from each group were trained to press a left lever for pellets and a right lever for a 20% sucrose solution, and the other half received the opposite arrangement. Because we have previously demonstrated that outcome devaluation performance is dependent on the dorsal hippocampus only during the early stages of learning, becoming hippocampally-independent after approximately 6 days of lever press training^43,44^, we first tested outcome devaluation after 4 days of training (“Day-4 Test”) so as not to miss this critical window. For this test, mice were given 1 hr of unrestricted access to one of the two outcomes so that they could consume it to satiety to reduce its value (devalued outcome) relative to the other outcome (valued outcome)^45^. This was immediately followed by a 10 minute choice test in which both levers were available but did not earn any outcomes. Mice that responded on the lever associated with the valued outcome and avoided the lever associated with the devalued outcome were considered goal-directed because they showed evidence of a) responding in accordance with outcome value, and b) responding in accordance with the lever press-outcome contingency as recalled from training, i.e. the two criteria of goal-directed action control^45^. Following this, mice received a further 4 days of training (8 days total) and were tested again (“Day-8 Test”), as we have previously found this to be sufficient for mice with hippocampal neuropathology to overcome deficits in devaluation testing^44^.

Figure 2B shows representative placements and spread of LPS based on the stereotaxic atlas of the mouse brain by Franklin and Paxinos^46^. Across Days 1-4 of lever press training, hippocampal neuroinflammation increased lever pressing in females but not in males (Fig. 2C). Statistically, although the main effect of sex was only marginal, F(1,32) = 3.96, p = .055, there was a main effect of LPS treatment, F(1,32) = 8.69, p = .006, and a 3-way sex x treatment x linear trend interaction, F(1,32) = 10.21, p = .003. This 3-way interaction was driven by a significant 2-way treatment x linear trend interaction for female mice, F(1,32) = 16.82, p < .001, but not male mice, F < 1, demonstrating that although the female LPS group increased responding faster than female Shams, male LPS animals did not differ from male Shams. Although the pattern of results could be interpreted as female Shams pressing at slower rates than all other groups, this is unlikely given that male mice are generally known to lever press at higher rates than females^44,47^. Rather, this result suggests that female LPS mice were pressing at abnormally high rates, on par with the baseline higher press rates in males.

On the Day-4 outcome devaluation test (Fig. 2D), neither male nor female Shams had yet acquired goal-directed control over their actions because they responded equally on Valued and Devalued levers. By contrast, male and female LPS-injected mice demonstrated intact outcome devaluation (Valued > Devalued), as supported by a devaluation x LPS interaction, F(1,32) = 6.679, p = .015, and main effects of devaluation, F(1,32) = 11.824, p = .002, and LPS treatment, F(1,32) = 4.642, p = .039, but not of sex, F < 1, and no 3-way interaction, F < 1.

Simple effects analyses confirmed that the devaluation x LPS treatment interaction reflected intact devaluation for female and male LPS groups, F(1,32) = 13.686, p = .001, for females, and F(1,32) = 5.486, p = .026, for males, with no significant effects in Shams of either sex, both Fs < 1. Importantly, these differences were not due to each group consuming a different amount of food in the prefeeding stage because consumption did not differ between groups, all Fs < 1 (Fig. 2E). This suggests that it is unlikely that group LPS did, but group Sham did not consume enough of each outcome for it to be devalued.

Mice next underwent an additional 4 days of lever press training (i.e. Days 5-8, Fig. 2F). During this training, we continued to observe sex-specific effects of hippocampal neuroinflammation on lever press responding because although there was no main effect of sex averaged across days, F < 1, there was a main effect of LPS treatment, F(1,32) = 8.252, p = .007, that interacted with sex, F(1,32) = 8.579, p = .006. Once again, this effect reflected greater responding in female LPS mice than in female Shams, F(1,32) = 15.542, p = < .001, with no such effect in males, F < 1. There was also a linear trend, F(1,32) = 261.66, p < .001, but in contrast to responding over Days 1-4 this did not interact with any other factor except sex, F(1,32) = 9.793, p = .004, all other Fs < 1. This suggested that all groups increased lever presses at the same rate, despite the differences in total presses. Overall, these results show that female LPS mice consistently responded more than female Shams on lever press training during Days 5-8, whereas male mice again performed similarly regardless of LPS treatment.

Following the last day of lever press training mice underwent another devaluation test, the “Day-8 Test” (Fig 2G). This time devaluation was intact for all groups, as supported by a main effect of devaluation (Valued > Devalued), F(1,32) = 26.822, p < .001, that did not interact with LPS treatment, F < 1, or sex, F(1,32) = 1.248, p = .272. There was also no 3-way interaction, F < 1, and no between-group main effects (largest F: main effect of Sex, F(1,32) = 2.379, p = .133). Once again, hippocampal neuroinflammation did not alter pre-test consumption of the outcomes (Fig. 2H), with the largest F(1,32) = 1.305, p = .262, for a main effect of sex. Therefore, in contrast to the Day-4 test, all groups exhibited goal-directed control on the Day-8 test, regardless of sex or LPS treatment.

Taken together, these results show that hippocampal neuroinflammation increases food-seeking actions (lever pressing) in female but not male mice, whereas it accelerates the acquisition of goal-directed action control in mice of both sexes, suggesting that it is sufficient to alter action selection and goal-directed action control in both consistent and sex-specific and ways.

### Hippocampal neuroinflammation impairs Pavlovian food-approach memories in females and facilitates them in male mice

As noted in the introduction, diseases that feature hippocampal neuroinflammation often affect food-related behaviours^16–18^ and do so in a sex-specific manner, with such impairments typically more prevalent in females^48–50^. Therefore, we next analysed whether hippocampal neuroinflammation was sufficient to produce these sex-specific disruptions to food approach responses. The instrumental testing and training protocols outlined above (see Fig. 2A) allowed us to measure this because they involve a Pavlovian food approach response wherein mice, in addition to performing the instrumental lever press, make head entries into the food magazine where food outcomes are delivered. Therefore, analysing these magazine entries provides a measure of Pavlovian food approach. Moreover, these protocols allowed us to determine whether food-approach itself is disrupted during training, when food is present, or whether the food-approach memories are disrupted during the devaluation test, which are conducted in extinction whereby food is absent.

Although lever pressing and magazine entry responses compete – it is (almost!) impossible for a mouse to do both simultaneously – we found that magazine entries were altered by hippocampal neuroinflammation in a manner that could not be attributed to competition with lever pressing. Rather, the differences between groups on each measure appear to have emerged independently.

First, in contrast to lever press responses during days 1-4 of acquisition which differed according to sex and LPS treatment, magazine entries during this period did not differ according to either factor (Fig. 3A), with no main effect of sex, F < 1, no main effect of LPS treatment, F(1,32) = 1.743, p = .196, and no sex × LPS interaction, F(1,32) = 3.02, p = .092. There was also no linear trend effect, F(1,32) = 3.022, p = .092, suggesting that magazine responding did not linearly increase or decrease across days.

**Figure 3:**
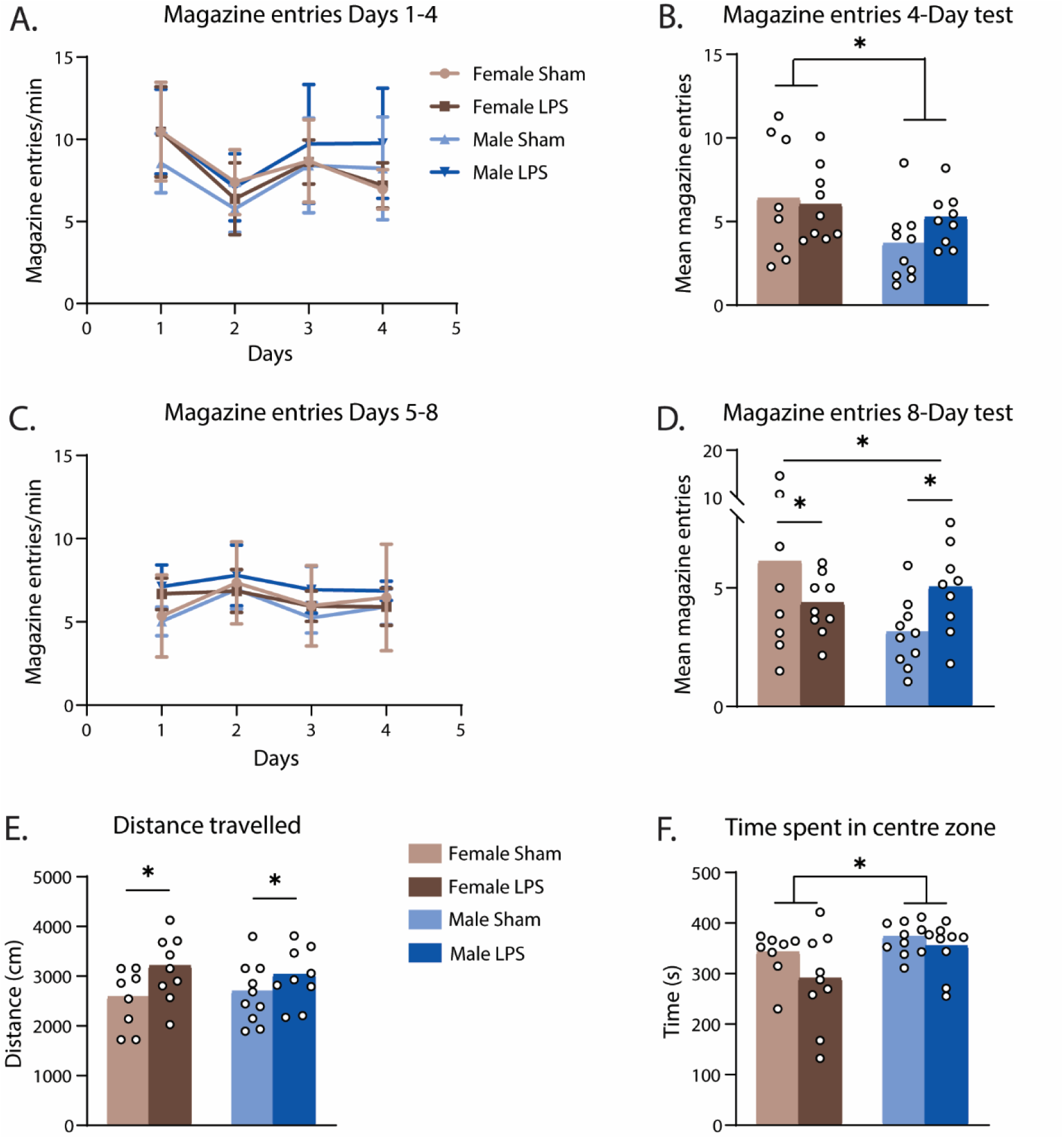
Hippocampal neuroinflammation has sex-dependent effects on Pavlovian food approach but increases locomotor activity for mice of both sexes. (A) Magazine entries per minute (±SEM) across days 1-4 of lever press training, (B) Mean magazine entries during the Day-4 test, (C) Magazine entries per minute (±SEM) across days 5-8 of lever press training, (D) Mean magazine entries during the Day-8 test, (E) Total distance travelled (F) and time spent in the centre zone during a 10 minute open filed test session for saline and LPS treated males and females. *p<0.05.

Magazine entries during the Day-4 devaluation test did differ according to sex, but not LPS treatment (Fig. 3B). Specifically, females made more magazine entries than males, but this did not interact with LPS treatment. This is indicated by a main effect of sex, F = 5.15, p = 0.03, no effect of LPS treatment, F < 1, and no sex x LPS interaction, F(1,32) = 1.353, p = .253. This result again differs from instrumental performance which varied in accordance with LPS treatment and not sex (i.e. the opposite pattern of results), suggesting that response competition cannot account for changes in either instrumental or Pavlovian responding. If it did, contrary to observations we would expect to see differences in accordance with the same factor in opposite directions, that is, increased magazine entries would lead to decreased lever pressing (and vice versa) in the same mice.

Magazine entries during days 5-8 of lever press training also did not differ between groups (Fig. 3C), as there was no main effect of sex, F < 1, no main effect of LPS treatment, F(1,32) = 3.027, p = .092, and no sex x LPS interaction, F(1,32) = 2.561, p = .119. Once again, there was no linear trend, F < 1, suggesting that magazine responding did not increase/decrease linearly across days.

On the Day-8 test, however, LPS treatment produced a doubly dissociable effect on magazine entries in females versus males (Fig. 3D). That is, whereas LPS treatment reduced magazine entries in females, it increased them in males. Statistically, there was no main effect of sex, F(1,32) = 1.914, p = .178, and no main effect of LPS treatment, F < 1, but there was a significant sex x LPS interaction, F(1,32) = 4.731, p = .037. Once again, this doubly dissociable effect does not appear related to magazine entries competing with the lever press response, because no group differences arose in instrumental responding on the Day-8 test.

These data show that although hippocampal neuroinflammation alters magazine approach behaviour during tests, when food is not delivered, it does not affect it during training when food is available. This suggests that hippocampal neuroinflammation specifically affects memories of food approach, rather than affecting food-approach behaviours directly. Moreover, just as in diseases featuring hippocampal neuroinflammation in which females display greater cognitive deficits than males^48,49^, here such neuroinflammation specifically impaired food-approach memories in female mice, whereas it strengthened them in males.

### Hippocampal neuroinflammation increased locomotor activity in mice of both sexes but did not affect anxiety-like behaviour in an open field test

We next questioned whether hippocampal neuroinflammation might have caused any broader alterations in locomotor activity or anxiety-like behaviours because again, both such behaviours have been shown to be altered in diseases featuring hippocampal neuroinflammation, often in a sex or gender-specific manner^19–21^. To this end, we conducted a 10 minute open field test to measure locomotor activity (the total distance travelled) and anxiety-like behaviour (the time spent in the centre zone versus the periphery). Typically, a more anxious mouse will avoid the centre and hide in the corners of the apparatus, akin to taking shelter from predators in the wild^51^, such that more time spent in the centre zone on this task is associated with less anxiety-like behaviour.

With regards to locomotor activity, LPS treatment increased the total distance travelled irrespective of sex (Fig. 3E), as indicated by a main effect of LPS treatment, F(1,32) = 5.562, p = .025, no main effect of sex, F < 1, and no sex x LPS interaction, F < 1. By contrast, anxiety-like behaviour differed according to sex but not LPS treatment (Fig. 3F). When time spent in the centre zone was analysed, there was a main effect of sex, F(1,32) = 5.56, p = .016, no main effect of LPS, F(1,32) = 3.081, p = .089, and no sex x LPS interaction, F < 1, showing that averaged across treatment groups, male mice spent more time in the centre zone than female mice, suggesting that female mice generally exhibited more anxiety-like behaviour.

Overall, these results show that hippocampal neuroinflammation increased locomotor activity but not anxiety-like behaviour in a manner that did not interact with sex, whereas anxiety-like behaviour was higher in female mice relative to males in a manner that did not interact with LPS treatment. These results suggest, therefore, that although hippocampal neuroinflammation is sufficient to disrupt locomotor activity, it is not likely to be the cause of any alterations to anxiety-like behaviours and is not the source of sex-specific alterations in either.

### Intra-hippocampal injections of lipopolysaccharide increased the expression of TNF-α, IBA1, and GFAP, and altered the morphology of cells positive for IBA1 and GFAP in the dorsal CA1 and dentate gyrus for mice of both sexes

Within one week of open field testing (approx. 1 month after stereotaxic surgeries) animals were transcardially perfused with PFA, brains were collected, and coronal sections containing dorsal hippocampus were immunostained for the pro-inflammatory cytokine tumour-necrosis factor-α (TNF-α), as well as microglial marker IBA1 and astrocytic marker GFAP, the increased expression of each of which has been considered to indicate neuroinflammation^52–54^. We here take the presence of IBA1 expression to indicate the presence of microglia, as it is specific to microglia in the brain^55^, although we note the possibility that some cells may be macrophages (a possibility that is still consistent with an inflammatory response).

The expression of TNF-α was quantified in two dorsal hippocampal regions that we identified as being those primarily targeted by our LPS injections: the dorsal CA1 and dentate gyrus. We chose to immunostain for this particular cytokine because not only does its presence indicate a neuroinflammatory response^54^, but increases in TNF-α have been linked to neuronal excitatabilty^56^. Thus, increased levels of this cytokine could provide a link between hippocampal neuroinflammation and any alterations in neuronal activation that might have occurred.

LPS injection led to the increased expression of TNF-α in both hippocampal regions, as expected (Results for CA1 shown in Fig. 4A-B, and for dentate gyrus results in Fig. 4C-D), and this difference was greater in females than in males. As TNF-α is not expressed in cell bodies, we limited our analysis to measuring its intensity (i.e. Mean Grey Value minus background). Unlike the cell culture study reported above, however, here we had two control (Sham) means and, as such, we here report the raw intensity numbers. In the CA1, there was a main effect of group (LPS > Sham), F(1,25) = 90.72, p < .001, that interacted with sex, F(1,25) = 8.63, p = .008, and this interaction was a result of a larger simple effect in females, F(1,25) = 61.73, p < .001, than in males, F(1,25) = 29.29, p < .001. Likewise in dentate gyrus, there was a main effect of group (LPS > Sham), F(1,25) = 127.46, p < .001, that interacted with sex, F(1,25) = 13.64, p = .001, which was again due to a larger effect in females, F(1,25) = 90.14, p < .001, than in males, F(1,25) = 38.23, p < .001.

**Figure 4:**
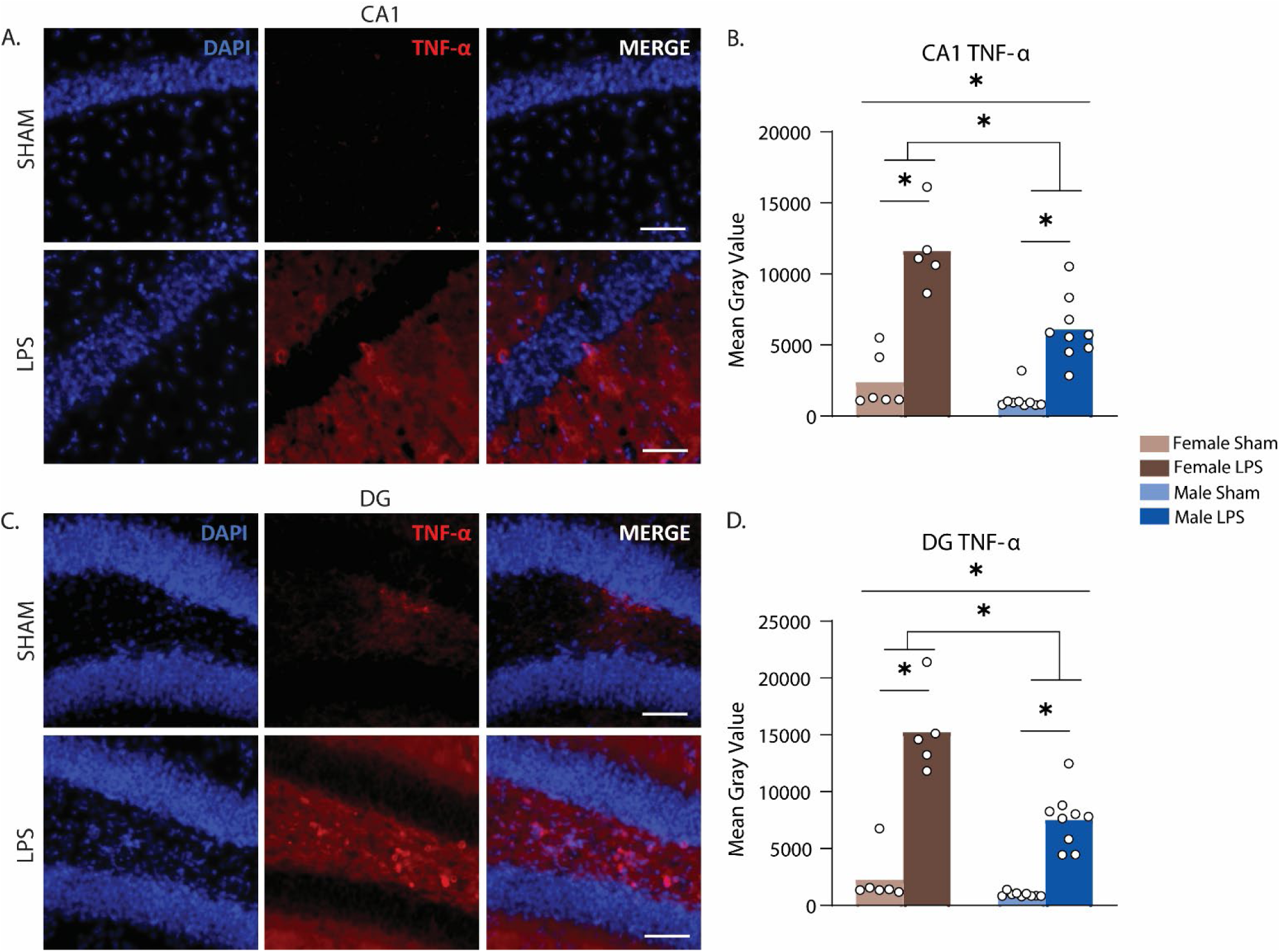
Intra-hippocampal lipopolysaccharide injections increased the expression of TNF-a in mice of both sexes, but to a greater extent in female mice. (A) Representative images of CA1 and (C) dentate gyrus regions of the dorsal hippocampus stained for DAPI-nucleus, tumour necrosis factor alpha (TNF-a; scale bar = 50µm) from a Sham (top panel) and LPS-injected mouse (bottom panel), (B) Mean Grey Value for TNF-a intensity in CA1, (D) Mean Grey Value for TNF-a intensity in dentate gyrus.*p<0.05.

Intra-hippocampal LPS injections similarly increased the intensity of IBA1 and GFAP in the CA1 and dentate gyrus of female and male mice, although this time the increases did not interact with sex (quantification of counts is reported in Supplemental Fig. 4). Figure 5A, from left to right, shows dorsal CA1 tissue, and Figure 5G shows dentate gyrus (DG) tissue, from a saline-injected Sham animal (top row) and an LPS-injected animal (bottom row). LPS increased IBA1 intensity in CA1 in both male and female mice (Fig. 5A), as indicated by a main effect of LPS, F(1,29) = 28.43, p < .001, no main effect of sex, F < 1, and no sex x LPS treatment interaction, F < 1. With regards to IBA1 intensity in DG (Fig. 5G), although the differences were directionally similar to that observed in CA1, they were not statistically significant, with no main effect of LPS treatment, F(1,29) = 2.181, p = .151, no main effect of sex, and no sex x treatment interaction, both Fs < 1.

**Figure 5:**
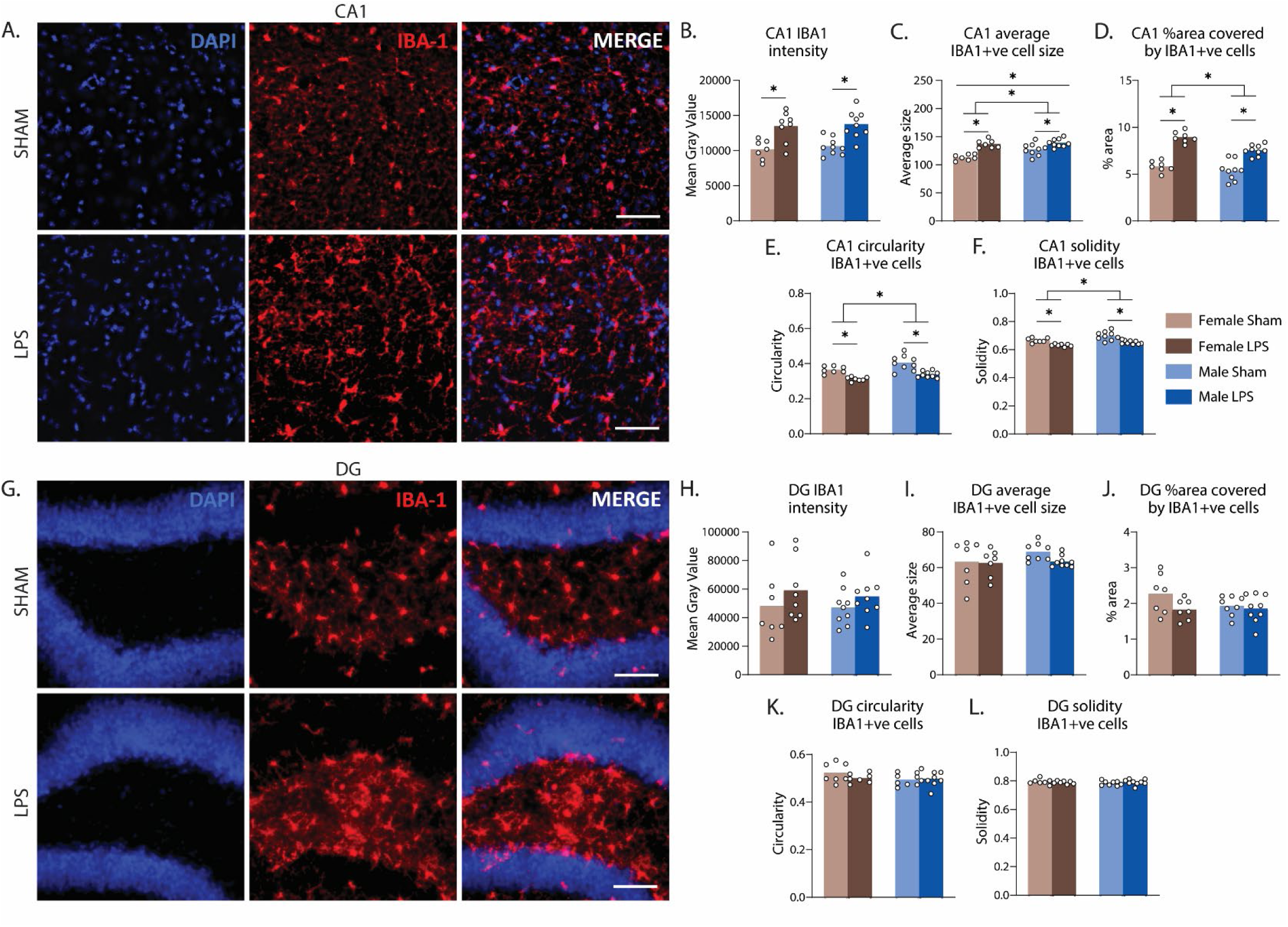
Intra-hippocampal lipopolysaccharide injections increased the intensity of IBA1 and altered the morphology of IBA1-positive cells in mice of both sexes. (A) Representative images of CA1 and (G) DG regions of the dorsal hippocampus stained for DAPI-nucleus, ionised calcium binding adaptor molecule (IBA1-microglia; scale bar-50µm), (B) Mean Grey Value for IBA1 intensity in CA1, (C) Average cell size of IBA1-positive cells in CA1, (D) Percentage of area covered by IBA1-positive cells in CA1, (E) Circularity of IBA1-positive cells in CA1, (F) Solidity of IBA1-positive cells in CA1, (H) Mean Grey Value for IBA1 intensity in dentate gyrus, (I) Average cell size of IBA1-positive cells in dentate gyrus, (J) Percentage of area covered by IBA1-positive cells in dentate gyrus, (K) Circularity of IBA1-positive cells in dentate gyrus, (L) Solidity of IBA1-positive cells in dentate gyrus. *p<0.05.

To capture more detailed morphological changes, we also quantified the average size of IBA1 and GFAP-positive cells, the total percentage of the area they occupied, their circularity, and solidity. We found that LPS led to a range of morphological changes in both cell types.

First in IBA1-positive cells in the CA1, LPS increased their average size, F(1, 28) = 37.69, p < .001 (Fig. 5C), and this effect was larger in females. There was an LPS x sex interaction, F(1,28) = 4.571, p = .041, supported by a larger simple effect for females, F(1,28) = 30.45, p < .001, than for males, F(1.28) = 9.157, p = .005. LPS also increased the percentage of area covered by IBA1-positive cells, F(1,28) = 98.15, p < .001, (Fig. 5D) although this did not interact with sex (F(1,28) = 3.13, p = .088). Both circularity (F(1,28) = 37.68, p < .001, Fig. 5E) and solidity (F(1,28) = 32.1, p < .001, Fig. 5F) were reduced by LPS, also in a manner that did not interact with sex (both Fs < 1). Similar morphological changes were not observed in the dentate gyrus (Figs. 5H-L, largest F was for % area, F(1,27) = 3.35, p = .078). These results demonstrate that, in CA1 at least, intra-hippocampal LPS injections increased the proliferation and altered the morphology of IBA1-positive cells.

Like with IBA1, LPS also increased GFAP intensity in both male and female mice in the CA1 (Fig. 6A). This is indicated by a main effect of LPS, F(1,29) = 67.86, p < .001, no main effect of sex, F < 1, and no sex x LPS treatment interaction, F(1,29) = 1.49, p = .215. This time, GFAP intensity was also increased by LPS treatment in both male and female mice in DG (Fig. 6B). Statistically, there was again a main effect of LPS treatment, F(1,28) = 38.57, p < .001, no main effect of sex, and no sex x treatment interaction, both Fs < 1.

**Figure 6:**
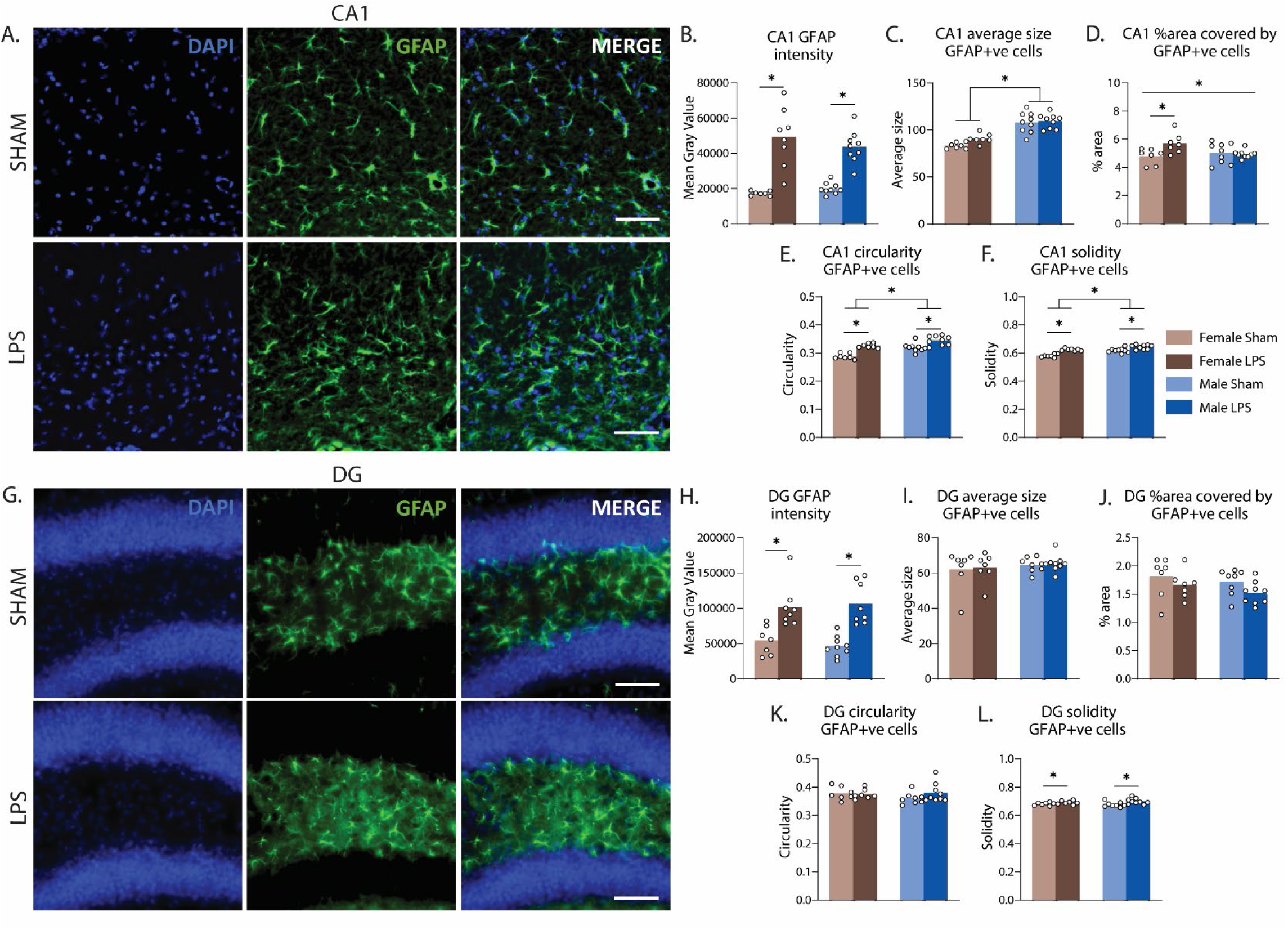
Intra-hippocampal lipopolysaccharide injections increased the intensity of GFAP and altered the morphology of GFAP-positive cells in mice of both sexes. (A) Representative images of CA1 and (G) DG regions of the dorsal hippocampus stained for DAPI-nucleus, glial fibrillary protein (GFAP-astrocytes; scale bar-50µm), (B) Mean Grey Value for GFAP intensity in CA1, (C) Average cell size of GFAP-positive cells in CA1, (D) Percentage of area covered by GFAP-positive cells in CA1, (E) Circularity of GFAP-positive cells in CA1, (F) Solidity of GFAP-positive cells in CA1, (H) Mean Grey Value for GFAP intensity in the dentate gyrus, (I) Average cell size of GFAP-positive cells in the dentate gyrus, (J) Percentage of area covered by GFAP-positive cells in the dentate gyrus, (K) Circularity of GFAP-positive cells in the dentate gyrus, (L) Solidity of GFAP-positive cells in the dentate gyrus. *p<0.05.

In CA1, LPS did not increase the average size of astrocytes, F(1, 28) = 2.715, p =.111 (Fig. 6E), but it did increase the percentage of the area covered by astrocytes, although only for females. There was no main effect of LPS, F(1,28) = 3.87, p = .059, but there was an LPS x sex interaction F(1,28) = 5.514, p = .026, which consisted of a significant simple effect for females (LPS > Sham), F(1,28) = 8.27, p = .008, and not for males, F < 1 (Fig. 6D).

LPS also increased the circularity of GFAP-positive cells (Fig. 6E), F(1,28) = 42.44, p < .001, in a manner that did not interact with sex, F(1,28) = 1.62, p .214, and it increased solidity (Fig. 6F) of GFAP-positive cells, F(1,28) = 46.89, p < .001, in a manner that also did not interact with sex, F(1,28) = 3.28, p = .081. Again, these changes were largely confined to the CA1 region, as there was no main effect of LPS on average size of astrocytes in the dentate gyrus, F < 1 (Fig. 6I), nor on the percentage of area covered by astrocytes, F(1,27) = 3.53, p = .071 (Fig. 6J), nor circularity (Fig. 6K), F < 1. There was, however, an increase in astrocyte solidity in the dentate gyrus of LPS-injected animals, F(1,27) = 7.5, p = .011, that did not interact with sex, F =1.23, p = .277, (Fig. 6L).

Together, these results confirm that LPS injected into the dorsal hippocampus was successful in inducing neuroinflammation in both male and female mice and provides some suggestion that the neuroinflammatory response may have been larger in female mice.

### Hippocampal neuroinflammation caused neuronal activation for female mice and reduced it for male mice

As mentioned previously, LPS-induced alterations in glial cells would be unable to effect behavioural changes due to the short processes of glia being unable to make contact with the broad neural circuit underlying motivated behaviour. Therefore, LPS must be exerting its effect by altering the activation of neurons, although our cell culture study indicates that this is not a direct alteration, but one that is mediated by glia. Our *in vivo* assessments revealed a range of LPS-induced alterations in glial marker intensity and morphology. Therefore, one final question to be answered is what changes in *in vivo* neuronal activation might have led to the observed behavioural changes. To answer this question, we investigated whether c-Fos expression was altered in neurons in the CA1 and dentate gyrus of the mice from our behavioural studies, and whether these changes were sex-specific.

We did this by immunostaining for the neuronal marker NeuN and the activity marker c-Fos in the same tissue, then calculating the percentage of c-Fos/NeuN co-localisation in the CA1 and DG regions of the dorsal hippocampus (individual counts of NeuN are included in Supplemental Fig. 4). Although we used MAP2 as a neuronal marker in our cell culture experiments in Figure 1, here we switched to NeuN because we found it to be a more reliable neuronal marker in fixed brain tissue, and easier to colocalise with c-Fos due to each being nuclear markers.

From left to right, Figure 7A shows tissue from the CA1 region taken from a saline-injected Sham animal on the top row, and an LPS-injected animal on the bottom row. Figure 7C shows tissue taken from the DG region, in the same order. The percentage area of colocalization of NeuN and c-Fos from the CA1 region is shown in Figure 7B, and the same measure from the DG region is shown in Figure 7D. In both regions, hippocampal neuroinflammation had a sexually dimorphic effect on neuronal activation: increasing it in females and decreasing it in males, although the evidence for this assertion is stronger in the DG. This is because, for CA1 sections, although there is a main effect of sex, F(1,28) = 7.422, p = .011, and no main effect of LPS treatment, F < 1, there was a borderline sex x treatment interaction, F(1,28) = 4.135, p = .052, although neither simple effect was significant for CA1 (largest F = 2.52, p = .124, for females (LPS > Sham)). In the DG, however, there was no main effect of sex or of LPS (both Fs < 1), but there was a significant sex x treatment interaction, F(1,27) = 20.185, p < .001, which, this time, was supported by a significant simple effect in females (LPS > Sham), F(1,28) = 12.19, p = .002, and a simple effect in the opposite direction for males (Sham > LPS), F(1,28) = 8.07, p = .009.

**Figure 7:**
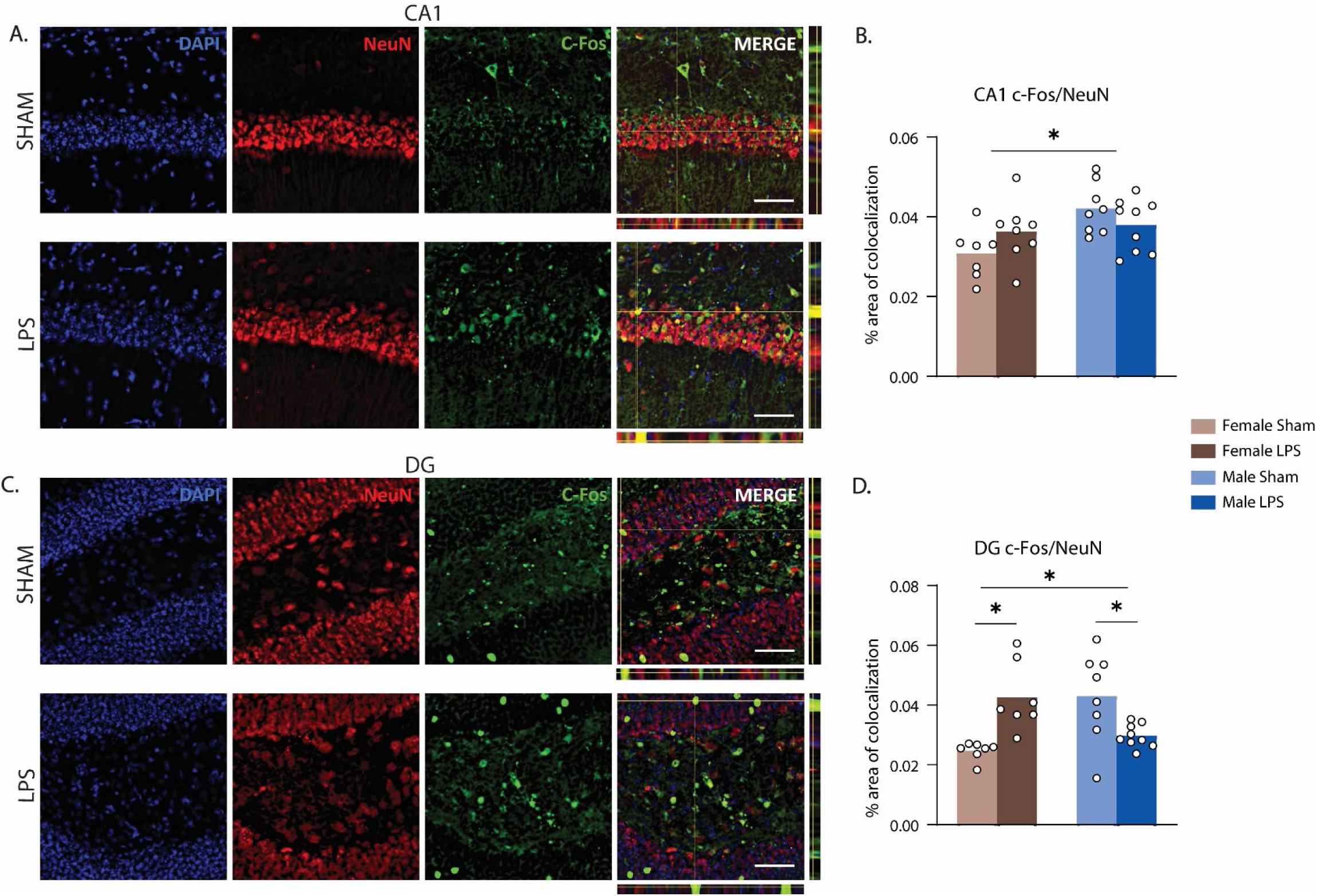
Intra-hippocampal lipopolysaccharide increased neuronal activation in females and decreased it in males in both dorsal CA1 and dentate gyrus. (A,C) Representative images of CA1 (A) and DG (C) regions of the dorsal hippocampus stained for DAPI-nucleus, NeuN-neurons, c- Fos-activated neurons (scale bar-50µm). (B) % area of co-localization of c-Fos and NeuN for CA1 and, DG region of dorsal hippocampus. *p<0.05.

These results suggest that hippocampal neuroinflammation does indeed produce sex-specific alterations in neuronal activation, increasing it in females yet decreasing it in males, particularly in the DG, suggesting this could be the potential underlying mechanism for the sex-specific behavioural differences observed.

## Discussion

Collectively, the results of this study show that hippocampal neuroinflammation is sufficient to produce sex-specific alterations in neuronal activation as well as several aspects of motivated behaviour. These findings suggest that hippocampal neuroinflammation could indeed be one underlying cause of disruptions to action selection, aspects of food-seeking and approach, and locomotor activity in diseases such as Alzheimer’s, multiple sclerosis, and depression.

First, we demonstrated that, in primary cell cultures containing hippocampal neurons alone, the application of an endotoxin known to trigger neuroinflammation had little effect, whereas applying it to primary cultures enriched with glia (astrocytes) caused neuronal activation as measured by an increase in c-Fos expression. This indicated a potential means by which neuroinflammation could ultimately affect behaviour. The first of our behavioural investigations revealed that hippocampal neuroinflammation selectively enhanced instrumental food-seeking actions (lever pressing) in female and not male mice but accelerated the acquisition of goal-directed control over actions in mice of both sexes.

Specifically, when tested following 4 days of lever press training, both male and female LPS-injected mice showed intact outcome devaluation (i.e. Valued > Devalued) compared to Sham controls (who responded Valued = Devalued). On the second devaluation test, conducted after 8 days of lever press training, all mice showed intact goal-directed performance (Valued > Devalued). Differences in satiety could not account for any of these effects because pre-test outcome consumption was comparable between all groups on both tests.

We next examined the effects of hippocampal neuroinflammation on Pavlovian food approach, as measured by head entries into the food magazine. Although we found no effects of sex or neuroinflammation on magazine entries during training, and a difference only in accordance with sex on the Day-4 test (females > males), we observed a double dissociation on the Day-8 test wherein LPS treatment reduced magazine entries in females and increased them in males. Again, this effect was unlikely to reflect an interaction with satiety, as consumption did not differ between groups. Moreover, the fact that this difference was observed only at test and not training suggests that hippocampal neuroinflammation had a specific and sexually dimorphic effect on memory for Pavlovian approach to a food-associated stimulus. With both general memory deficits and specific food-seeking deficits prevalent in diseases such as Alzheimer’s^16,19^ and depression^17^, each of which have been argued to be more severe in females^48,57,58^, this finding suggests that hippocampal neuroinflammation could be the underlying cause of these sex-specific cognitive impairments. Our final behavioural assessment using the open field test revealed that hippocampal neuroinflammation increased locomotor activity in mice of both sexes but did not alter anxiety-like behaviour, although this was higher in females in a manner that did not interact with LPS treatment: an effect that is variably reported in the literature^47^.

Interestingly, our findings are at odds with several reports of systemic administration of LPS reducing locomotor activity, food intake, and impairing cognition^59–61^. It is possible that these differences between studies are due to peripheral LPS causing sickness behaviours, whereas our intra-hippocampal administrations of LPS caused minimal or no sickness behaviours (see methods for details).

Our immunohistochemical analyses showed that LPS injections were effective in producing neuroinflammation in dorsal CA1 and dentate gyrus, because we observed increased intensity of TNF-α, IBA1 (microglial), and GFAP (astrocytic) expression in LPS-treated animals, as well as several morphological changes in both cell types. It is worth noting that not all these alterations were in the expected direction, however. In particular, the findings that LPS increased cell size and decreased circularity in IBA1-positive cells were at odds with several^53,62–64^ prior studies (although they are consistent with others^65^). These discrepancies may stem from differences in study time lines or brain regions, as prior studies examined microglial morphology hours or days rather than 1 month post-insult, and many of them studied microglia from outside of the hippocampus where morphology can vary considerably^28,63,66^. Nevertheless, LPS clearly induced changes in cell proliferation and morphology in both male and female mice, with some differences more pronounced in females. This neuroinflammatory response had a sexually dimorphic effect on neuronal activation, increasing it in females and decreasing it in males, providing potential insight into the underlying mechanism of the sex-specific behavioural effects.

### The behavioural disruptions produced by hippocampal neuroinflammation are predominantly independent of one another

Most behaviours investigated here are underpinned by motivation, emotion, and motor control, suggesting that the disruption of just one of these underlying constructs could have led to the myriad disruptions observed. However, a close look at the observed pattern of behavioural disruptions does not reveal one singular factor that could account for all the behavioural changes. For example, the general increase in locomotor activity could be assumed to account for the increase in lever pressing and the acceleration of goal-directed action control in female LPS mice. That is, if LPS-injected females simply moved more than Sham controls, this could account for the increase in lever presses. More lever presses, in turn, meant that the female mice received more lever press-outcome pairings which could have led to the accelerated acquisition of goal-directed control. Such an account cannot be equally applied to the LPS-injected male mice, however, who also demonstrated an increase in locomotor activity and the acceleration of goal-directed control, but who pressed the levers at similar rates to male Shams and thus received the same number of lever press-outcome pairings. This suggests, therefore, that the behavioural effects are independent of each other, at least in males. Although it is possible that male Shams lever press rates were at ceiling, precluding similar observations of elevated pressing as seen in female LPS mice, this seems unlikely given that mice are capable of lever pressing at much higher rates than we observed here^67,68^.

The behavioural changes in instrumental and Pavlovian responding likewise appear to be independent, despite lever presses and magazine entries being concurrently available throughout most behavioural training and thus having the potential to compete. As detailed in the Results, response competition would have meant that an increase in lever pressing should have led to decreased head entries into the magazine, and vice versa, as mice typically cannot perform both actions at the same time. However, such an inverse relationship was not observed during any period of training or test, suggesting that the various effects are unrelated.

Furthermore, none of the behavioural changes caused by neuroinflammation are likely to have resulted from anxiety-like behaviours, because this was unaffected by LPS treatment and differed only in accordance with sex. It is conceivable that neuroinflammation interacted with the higher anxiety in female mice to produce the sex-specific differences observed in instrumental responding and goal-directed action control, although if this were the case it would once again be more likely to impair rather than facilitate and accelerate these behaviours. This is because these kinds of appetitive and aversive behaviours are commonly assumed to be mutually inhibitory^69,70^.

Overall, therefore, the most parsimonious account of the current results is that hippocampal neuroinflammation produces a broad array of alterations to motivated behaviour, in a manner that is sometimes dependent on sex, much as it is proposed to do in humans^2,7,9–11,19^.

### The acceleration in goal-directed action control does not indicate that hippocampal neuroinflammation improves cognition

The direction of several of the differences between animals with hippocampal neuroinflammation and controls could be interpreted as evidence that neuroinflammation improved rather than impaired learning, a finding that would be surprising given the noted cognitive-behavioural impairments in individuals with diseases featuring such neuroinflammation^7,10,11^. A notable example is the accelerated acquisition of goal-directed action. However, we suggest caution in interpreting this as evidence that neuroinflammation “improved” goal-directed control. Indeed, when a similar result was demonstrated in the K3691 tau mouse model of Alzheimer’s disease^71^ the authors suggested that, rather than improving behaviour, the transgenic manipulation had impaired development of habits relative to controls. A similar interpretation could be applied to current results, because habitual responding on devaluation tests produces performance that is identical to that exhibited by male and female shams on the Day-4 test in the current study (i.e. Valued = Devalued). This interpretation seems less likely, however, when it is considered that the same Sham controls in this study underwent additional lever press training, after which they displayed intact devaluation on the Day-8 test (i.e. Valued > Devalued). If Shams were habitual on the Day-4 test, the additional days of training would be expected to strengthen the habit such that the equivalency of responding on each lever should continue to be observed on the Day-8 Test, but this this was not the case. Therefore, we can rule out this interpretation.

Nevertheless, the results of the Day-4 test do provide evidence that hippocampal neuroinflammation disrupts the balance between habitual and goal-directed actions. That is, although behaviour exhibited at any one time is thought to be under the control of one system or the other, both goal-directed and habitual *learning* is present from the very beginning of training. The behavioural output on any test simply reflects the summation of this learning to reveal which system is currently more dominant^36,45,72^. On this view, if the performance of Sham animals is taken to represent the optimal balance between goal-directed and habitual control at the time of the Day-4 test, then it could be assumed that hippocampal neuroinflammation altered this balance in a manner that is sub-optimal.

Indeed, excessive goal-directed control has been identified as problematic in several diseases and disorders that feature hippocampal neuroinflammation^73,74^, and in the many day-to-day situations where executing fast, automated habit-based responses is optimal, persistent or premature goal-directed responding is likely maladaptive. For example, being “hyper goal-directed” has been suggested to contribute to the slowness of actions in individuals with Parkinson’s^75^, or to drug-seeking in substance use disorder at the expense of other, healthier choices by powerfully elevating drug value^73^. The increases in locomotor activity and instrumental responding could be considered in much the same way. Both increases and decreases in motivated behaviour have been observed across most of the diseases involving hippocampal neuroinflammation^10,11,16,20,21^, and changes in either direction can be considered maladaptive.

A final possibility is that, in females at least, the neuronal activation caused by neuroinflammation did in fact improve learning but that such an effect is limited to the time frame captured here (i.e. with devaluation testing occurring 2-3 weeks after LPS administration). However, the persistence of neuronal activation that occurs in response to an insult rather than to explicit changes in environmental stimuli has been suggested to signal the subsequent death of those neurons^76^ (but see^77^), which could then impair the same behaviours that were initially facilitated. If this were the case, it would explain why our previous study of outcome devaluation in a J20 mouse model of Alzheimer’s disease found goal-directed learning to be impaired relative to wildtype controls^44^, despite similar levels of hippocampal neuroinflammation to that observed here. Age might also matter here: our mice were 9-10 weeks old at the start of each experiment, broadly corresponding to the end of adolescence and start of young adulthood in mice^78^. Because immunoreactivity of brain cells alters with age^79,80^, future studies may therefore wish to repeat the current study in older animals to determine whether the pattern of behavioural changes persists or is altered.

### Are the effects of neuroinflammation effects on neuronal activation and behaviour specifically mediated by astrocytes

The results of the cell culture study indicate that the differences in *in vivo* neuronal activation are unlikely to have occurred via a direct effect of the LPS injections on neurons, possibly because LPS is a toll-like 4 receptor agonist, and these receptors are expressed more densely on glia than on neurons^81^. Rather, the differences in neuronal activation likely occurred due to modulation by glia, and particularly astrocytes. Therefore, given that LPS increased the quantification of GFAP signal intensity in both male and female mice, it is somewhat surprising that neuronal activation was only observed in the post-mortem immunohistochemical investigation in female mice, and was reduced in male mice. One possible reason for this difference could be the different time-scales in the two studies: in our cell culture study, the effects of LPS on neuronal c-Fos were investigated 1 day following application, whereas for our *in vivo* study, the effects were not evaluated until 1 month later. Therefore, it is possible that these differences led to an acute versus a chronic neuroinflammatory response, which had differential effects on neuronal activation. However, why this would differentially affect neuronal c-Fos in males versus females is unclear.

As shown in Supplemental Figure 4C and 4D, when averaged across treatment groups we observed a higher number of astrocytes in the dorsal hippocampi of females relative to males, which perhaps created more opportunity for neuronal activation upon the administration of LPS. Even if this were the case, however, it is not clear why neuronal activation was reduced in LPS-injected males. Because the cells for our cell culture study were extracted from animals that were not sexed (as it was not determined that sex influenced neuronal activation in response to LPS administration until after cell culture experiments were complete), it is possible that they were predominantly sourced from female animals, and this is the reason for the observation of neuronal activation rather than inhibition. Either way, the precise mechanisms of astrocytic modulation of neuronal activation is unclear from the current findings and warrants further investigation.

Future studies might particularly wish to investigate whether the activation of astrocytes alone is sufficient to replicate the changes in motivated behaviour. We did already make one attempt to answer this question using astrocyte-specific chemogenetics, specifically hM3Dq designer receptors exclusively activated by designer drugs (DREADDs) under the GFAP promoter. However, despite being well powered (post placement samples sizes: n = 33 females and n = 32 males), this study did not yield any significant behavioural results of interest, possibly due to additional stress caused by the additional CNO/Vehicle intraperitoneal injections that were given prior to each training session (for any interested researchers the full dataset is available at the same link as the data for the current study – see methods). These results were further confounded by the fact that DREADD transfection was not astrocyte-specific because we identified partial colocalisation with NeuN. Therefore, future studies addressing this question may want to explore the use of different methodologies.

### Hippocampal neuroinflammation and sex hormones

Another key question for future studies is the mechanism by which the same manipulation – LPS injections – produced different behavioural outcomes and differential neuronal activation in male and female mice. Sex hormones comprise a plausible candidate because the presence of estrogen has been shown to increase neuronal excitability within the hippocampus^82,83^, whereas androgens appear to decrease it^84,85^. Although these studies were conducted in the absence of neuroinflammation, sex hormones and particularly estradiol are seen as neuroprotective^86,87^, for instance reducing the number of pro-inflammatory cytokines in the brain following LPS injection^88^. Levels of estrogen have also been reported to increase following neuroinflammation in both males and females, to facilitate brain repair^89,90^. Thus, it is possible that an interaction between microglial/astrocytic responses and estrogen (or androgens) occurred in the current study, particularly in females, that could account for the changes in behaviour and neuronal activation.

## Conclusion

Here we have presented the first demonstration, to our knowledge, that hippocampal neuroinflammation is sufficient to disrupt both neuronal activation and multiple aspects of motivated behaviour in either consistent or sex-specific ways. This is important because although numerous studies have demonstrated sex differences in glial and immune responses^55–58,91^ as well as in behaviour^92,93^ the current study is the first to connect the two in a causal manner. Moreover, prior to the current study it was unclear whether hippocampal neuroinflammation, which is common to many diseases and disorders, was simply another symptom or whether it could alter these behaviours in the absence of additional neuropathologies. We have shown that it is the latter. Although not addressed in the current study, it would be of interest for future research to assess the broader neural circuit that is affected by hippocampal neuroinflammation to produce the observed effects. For instance, prelimbic cortex^94,95^ and dorsomedial striatum^96,97^ have been heavily implicated in the mediation of goal-directed action control, and it would be of interest to determine whether hippocampal neuroinflammation alters neural activity in these regions to achieve the effects observed here. Finally, although current findings do not suggest that hippocampal neuroinflammation is solely responsible for the behavioural changes observed in each of these diseases, they do suggest that it is likely to contribute – at times in a sex-specific manner - and therefore targeting hippocampal neuroinflammation, particularly in females constitutes a promising therapeutic objective for preventing these behavioural changes.

## Acknowledgements

This work was supported by the National Health and Medical Research Council of Australia Grant GNT2003346 and by the Australian Research Council Grant DP200102445 to L.A.B. We thank Genevra Hart, Melissa J. Sharpe, Claire I. Richards, and Bernard W. Balleine for helpful discussions regarding this manuscript. We thank the technical staff at the Garvan Biological Testing Facility at the Garvan Institute of Medical Research and the technical staff at the Ernst Facility at the University of Technology Sydney for technical support. Figures 1A and 2A were created with <u>BioRender.com</u>.

## Author contributions

K.G., S.G., S.T.B., S.B., and L.A.B., conceptualised and designed the research, K.G., S.G., W.K., S.T.B., J.M.G., A.D., A.A., M.M., A.C., M.K., V.V., and L.C., performed the research (i.e. data curation), K.G., S.G., W.K., J.M.G., and L.A.B., analysed the data, L.A.B., acquired the funding, K.G., S.G., W.K., J.M.G., M.K., A.C., and L.A.B. wrote the paper (original draft – K.G., S.G., W.K., and L.A.B., review and editing - K.G., S.G., W.K., J.M.G., M.K., A.C., and L.A.B.).

## Data and code availability

Data: Behavioural data have been deposited at Open Science Framework and are publicly available as of the date of publication. Microscopy data reported in this paper will be shared by the lead contact upon request.

Code: This paper does not report original code.

Any additional information required to reanalyse the data reported in this paper is available from the lead contact upon request.

## EXPERIMENTAL MODEL AND SUBJECT DETAILS

### Primary Cultures

Primary mouse astrocytes and microglia were isolated from SwissTacAusb mice at P0-P2. We did not determine the sex of mice at this age as it is very difficult to determine from physical inspection, and excessive handling causes stress to the animals. Mouse brains were extracted, and hippocampi isolated and dissociated by incubating with 2.5% trypsin/EDTA. Samples were centrifuged and cell pellets were triturated to generate single cell suspensions to allow seeding on T75 flasks pre-coated with 0.1% poly-D-lysine (PDL) at 1-1.5 x10^6^ cells/mL in warmed culture media consisting of DMEM/F12 with 10% FBS and 1% P/S. Flasks were incubated at 37°C in 5% CO_2_, and culture media was changed every 2-3 days for 10 days, when flasks were oscillated at 200rpm for 2.5 hours to separate microglia and astrocyte layers. The supernatant was collected and centrifuged to obtain enriched microglia pellet, while the remaining astrocytes were incubated with 2.5% trypsin/EDTA before being collected and centrifuged to obtain enriched astrocyte pellet. Microglia and astrocyte cell pellets were seeded on separate PDL coated flasks in culture media at 1-1.5 x 10^5^ cells/mL and 2.4-2.7 x 10^5^ cells/mL respectively, reaching confluency after 2-3 days.

Primary mouse cortex and hippocampal neurons were purchased from Sigma Aldrich. Because these cells were obtained independently (by Sigma Aldrich) so the sex and age of animals is unknown. Vials were rapidly thawed (from -130C) and contents transferred to conical culture tubes in warm Neurobasal medium with 0.5 mM GlutaMAX and 2% B-27 (ThermoFisher Scientific). 10µL was removed and added to a microcentrifuge tube with 0.4% trypan blue for a viable cell count. Cells were then seeded at 4-8 x10^5^ cells/ mL onto PDL coated flasks in Neurobasal medium and incubated at 37°C in 5% CO_2_. Half the volume of media was replaced every third day until cells reached confluency after 10 days, monitoring all cultures daily to track growth and discern contamination.

### In vivo Animal Studies

C57BL/6J healthy and experimentally-naïve mice aged 9-10 weeks at the beginning of the experiments were housed at a maximum of 5 in a cage throughout the experiment. All mice were maintained on a 12-hour light/dark cycle and had ad-libitum access to water ad-libitum. All animals weighing a minimum of 25gm were obtained from Australian Bioresources facility where the animals have been backcrossed for multiple generations. Lights were turned on at 7am and all behavioural experimentation occurred during the light cycle. Room temperature was kept at a consistent 21-22 degrees celsius. A total of 22 females 24 males were used in the experiment. 5 out of 22 females and 4 out of 24 male animals were excluded from behaviour analysis because of incorrect needle placement. 1 male animal was excluded from the data due to being a consistent statistical outlier, performing two standard deviations above the mean. Apart from the above-mentioned animals, an additional 2 females and males were removed from the immunoanalysis due to tissue damage that occurred post-mortem. Irradiated chow was provided from specialty feeds and was provided ad-libitum except when animals were food deprived 3 days prior to, and during experimental testing. During food deprivation animals received 1.2-1.3g per mouse (1.2g per female, 1.3g per male) per day. Female and male mice were randomly assigned to saline and LPS treatment groups. All animal experiments were performed with the approval of the Garvan Animal Ethics Committee and University of Technology Animal Care and Ethics committee, and the guidelines set out by the American Psychological Association for the treatment of animals in research. Specific details of mice in each experiment are provided below.

## METHOD DETAILS

### Cell culture study Methods

#### Culture conditions and models of inflammation

The following experimental settings were employed: (i) individual monocultures of neurons, astrocytes, and microglia, (ii) cocultures of neurons-astrocytes and neurons-microglia, (iii) tricultures of neurons, astrocytes, and microglia. Co- and tricultures were established using monocultured cells at proportions of 2 neurons to 5 astrocytes and 1 microglia. All cultures were seeded on PDL coated flasks and incubated in a 37°C humidified atmosphere with 5% CO2, monocultured neurons in Neurobasal culture medium with 0.5mM GlutaMAX and 2% B-27 (Thermofisher Scientific), monocultured glial cells in DMEM/F12 (Sigma) with 10% FBS and 1% P/S, and co- and tri- cultures in 70% neuron medium and 30% glial cell medium. We repeated experimental settings with cells from three biological replicates. Cultures were treated with 1 μg/mL LPS for 24 hours, with untreated cultures as controls.

#### Immunofluorescence staining

Following the experimental treatment, cultures were washed with phosphate buffered saline (PBS) and fixed with 4% w/v paraformaldehyde (PFA; Sigma) with PBS for 1 hour. Fixed cells were washed with 0.02% v/v Tween20 (Sigma) in PBS, and permeabilised with 0.1% v/v Triton X-100 (Sigma) in PBS solution for 3 minutes. A solution of PBS with 5% v/v goat serum (Invitrogen) and 3% BSA was added as a blocking buffer, for 3 hours. Samples were incubated overnight in a blocking buffer and primary antibody solutions containing the following: mouse anti-IBA1 (Invitrogen, 1:250), rabbit anti-IBA1 (Abcam, 1:250), mouse anti-GFAP (Cell Signalling Technology, 1:150), chicken anti-MAP2 (Abcam, 1:500), and/or rabbit anti-c-Fos (Synaptic Systems, 1:500). After incubation, cells were washed with PBS, then incubated for 1 hour in a solution of PBS and secondary antibodies; DAPI (Invitrogen, 1:1000), goat anti-rabbit-AlexaFluor 488 (Abcam, 1:1000), goat anti-chicken-AlexaFluor 555 (Abcam, 1:1000) and/or goat anti-mouse-AlexaFluor 647 (Abcam, 1:1000).

#### Microscopy

*In vitro* images were acquired with the TiE2 widefield fluorescence and transmitted light microscope with Andor Sona camera (MIF, UTS), with both 10X objective (NA=0.45), WD=4000µm) using 2.5µm optical slices, and 20X objective (NA=0.70, WD=2300µm) with 1µm optical slices. Analysis was conducted on 20X images, 5 images per control/LPS, 10 images per culture condition, 6 culture conditions per biological replicate for a total of 120 images for 12 cultures.

#### Cytokine Assay

A panel of cytokines was measured in the supernatant of the neuron/astrocyte/microglia tricultures using a mouse inflammation antibody array membrane (Abcam), according to the manufacturer’s instructions. Briefly, membranes were incubated with blocking buffer for 30 minutes at room temperature. Membranes were probed with 1 ml of cell supernatant of the tricultures for overnight at 4°C , and then washed and incubated 1 ml with Biotin-Conjugated Anti-Cytokines. Membranes were again washed, incubated with 2 ml of HRP-Conjugated Streptavidin and incubated overnight at 4°C. Finally, membranes were washed and incubated with 500 μl of the Detection Buffers mixture for 2 min at room temperature. Membranes were imaged using a ChemiDoc MP Imaging System (Bio-Rad) for exposures of 3 min and images were visualized and analysed using a protein array analyser plugin in FIJI.

### Antibodies

Iba1 ms invitro (1:250)

Iba1 rb abcam (1:250)

c-Fos ms abcam (1:500)

rabbit pAb-c-Fos (synaptic systems)

GFAP – ms cell signalling (1:150)

MAP2 – chk abcam (1:500)

AF594 anti-rabbit (1:1000)

AF647 anti-chicken (1:1000)

Af488 anti mouse (1:1000)

New c-Fos – 1:200 (abcam)

PDL coating – 4.5 µg/cm^2^ – made by adding 5mg PDL to 50mL tissue culture grade water.

### *In Vivo* study Methods

### Drugs

Lipopolysaccharide (LPS, *Escherichia coli* O111:B4, Sigma Aldrich) was injected intra- hippocampally at a concentration of 4µg/µl. A Ketamine (100 mg/kg of body weight, Mavlab) + Xylazil (10mg/kg of body weight, Troy laboratories Pty ltd) mixture was used for stereotaxic surgery and cardiac perfusion procedures. All drugs were diluted to working concentrations in Saline (0.9% Sodium chloride injection BP, Pfizer).

### Stereotactic surgery

Mice were anaesthetized via intraperitoneal (i.p.) injection with the ketamine + xylazil mixture noted above and then fixed on a stereotaxic frame (Model 940, David KOPF Instruments). An incision of approx. 2cm was made on the skin with a scalpel blade (size 22) to expose the skull of the animal. The brain was levelled, and small holes were drilled (microdrill, SDR scientific, Harvard Apparatus) on either side of the skull at the co-ordinates (in mm, from bregma): anterior-posterior: -1.8mm, mediolateral: +/-1.5, dorsoventral, - 1.7mm. A Hamilton syringe (10µl, 1700 series, RN syringe, 1.0µl, 7000 series, KH syringe, Neuros syringes, SDR scientific, Harvard Apparatus) filled with LPS/Saline was used to inject the desired solution (1µl per hemisphere) at the rate of 0.2μl/minute. The syringe was then left undisturbed for 5-7 minutes post injection and slowly retracted to make sure the liquid was contained in the area of injection. All animals were given Carprofen (Rimadyl, Zoetis) 0.05ml and Bupivacaine (Sterisafe, Pfizer) 0.1ml during surgery to aid a faster recovery. The wound was then sutured (Dysilk, S405, 18mm, 3/8 circle, Dynek Pty Ltd). Mice were closely monitored during recovery for signs of weight loss, pilo erection, hunched body posture, inflammation, gripping strength, and any infection around the surgical site. These were recorded in the monitoring sheet with a grimace scale rating of 0-2. All but two animals scored zero after the surgery (i.e. no grimace present). These two animals scored 1 for two days post surgery which then shifted to zero and remained at zero for the rest of the experiment. All animals were weighed and monitored daily for a minimum of 7 days while recovering from surgery. Body weights remained above 95% of initial weights during post-surgery recovery, prior to food deprivation.

### Apparatus

The appetitive instrumental portion of the experiment was conducted in 6 operant chambers (Med Associates). One side of the chamber wall consisted of a magazine (food receptacle) in the center and two retractable levers one on either side of the magazine earned either a 25 mg grain food pellet (Bioserve Biotechnologies) delivered to the left well of the magazine via a pellet dispenser, or 2mL of a sucrose solution (20% white sugar, Woolworths, Australia, and 10% maltodextrin, Poly-Joule, Nutrica, Australia diluted in H2O) delivered to the right well of the magazine via syringe pump. The opposite side wall consisted of a house light for illumination and a house fan which provided constant ∼70 dB background noise, both of which were present throughout all behavioural procedures unless otherwise stated. All training sessions were pre-programmed on two computers through the MED Associates software (Med-PC), which also recorded the experimental data (lever presses, outcome deliveries, magazine entries) from each session.

The locomotor and anxiety assessments took place in four open field chambers that measured 273mm x 273mm with 203mm high glass walls. These were placed inside a sound attenuating cubicle (MED-OFAS-MSU, MED-OFA-022, Med Associates inc.). Movement was recorded and tracked with Activity Monitor 7 (Med Associates inc.) which uses Infrared beams.

### Behavioural Procedures

#### Food Deprivation

Three days before the start of magazine training, mice were food deprived with male mice receiving 1.3g chow and female mice receiving 1.2g chow. The food deprivation continued for the duration of behavioural procedures; however, the quantity of chow was increased after 2 weeks to 1.6g for males and 1.4g for females to prevent body weights failing below 80% of their pre-surgery body weight. Weights were monitored 3 times a week on Mondays, Wednesdays, and Fridays.

#### Magazine pre-training

Mice received magazine training for the first two days to familiarize them with the operant chambers and reduce neophobia. During this session, sucrose and pellet outcomes were delivered to the magazine at random intervals around a mean of 60s. The session terminated after 30 minutes or when 20 sucrose and pellets (40 total outcomes) had been delivered, whichever came first. Levers were not present during magazine training.

#### Lever-press training (Day 1-4)

Lever press training commenced one day following magazine pre-training. For half of the animals in each group, the left lever earned sucrose and the right lever earned pellets, and the rest of the animals received the opposite arrangement (counterbalanced, randomly assigned). Each lever press session consisted of four 10 minute periods during which a single lever was extended and earned a single outcome (i.e., 2 x 10 minute per lever), the order of which was randomized each day. Each 10 minute sub-session was separated by a 2.5 minute period during which both levers were retracted, and the house lights turned off. Sub-sessions terminated and mice entered the 2.5 minute interval early if 20 outcomes were earned before 10 minutes had elapsed. All mice were trained on a continuous reinforcement schedule (CRF, each lever press earns a food outcome) for Days 1-2 and were increased to random ratio schedules 5 (RR5: i.e. each lever press earned an outcome with a probability of 0.2) on Day 3, then to RR10 (i.e. each lever press earned an outcome with a probability of 0.1) on Day 4. Lever press sessions terminated after 50 minutes or when animals had earned 40 of each outcome, whichever came first.

*Day-4 outcome devaluation Test:* All mice received a 1 hr pre-exposure session to the devaluation boxes containing a small portion of their daily chow allocation following the last lever press training session on Day 4 to reduce neophobia. The next day, the first round of outcome devaluation testing was conducted, collectively termed “the Day-4 test”. Mice were given 50 minutes of ad libitum access to either sucrose or pellets (randomly assigned, counterbalanced) to reduce the value of the prefed outcome relative to the other outcome (Balleine & Dickinson, 1998). Mice were then immediately placed into the operant chambers for a 10 minute devaluation test where they were presented with both levers available, but no outcomes were delivered (i.e., testing occurred in extinction). The following day, mice were tested again in an identical fashion except that they were prefed with the opposite outcome. All test data is reported collapsed across the two days of testing.

*Lever press training (Day 5-8)*: The day after the Day-4 test, animals were re-trained to lever press in the manner previously described. Mice were trained on an RR5 schedule on Day 5, then on an RR10 schedule for days 6-8.

*Day-8 outcome devaluation test:* The day following Day 8 of lever press training mice were given a second round of outcome devaluation testing, referred to as the ‘Day-8 test’. This was conducted identically to the 4 Day test described above.

#### Open Field Test

Individual mice were placed in the centre of the chamber and given 10 minutes to explore. Distance travelled and time spent in the centre zone of the chamber versus the corner zone was tracked and measured by Activity monitor 7 (Med Associates software).

### Tissue collection

Mice were anaesthetized with the Ketamine + Xylazil mixture mentioned above perfused 15 minutes later. Mice were cut open from the abdominal region till the rib cage to reveal the heart. A 27G needle was used to puncture the apex of the heart and an incision was made in the right atrium. 0.9% Saline was first delivered for one minute to flush the blood from blood vessels. Mice were then perfused with ice-cold 4% paraformaldehyde (PFA) in phosphate buffer saline (PBS, at pH 7.4) for approx. 8 minutes. Brains were carefully extracted and stored in 4% PFA overnight. The next day, brains were moved to a 30% PBS sucrose solution where they were stored until sectioning.

Brain sections of 40μm thickness were cut using cryostat with an interval of 1:6 and stored in cryoprotectant (0.2M Phosphate buffe, Ethylene glycol, glycerol and milliQ) solution. Sections were then rinsed with PBS 3 x 10 minutes and blocked with 3% BSA (Bovogen Biologicals, BSAS 1.0) + 0.25% Triton (Sigma Aldrich) in 1x PBS (pH 7.2) for an hour at room temperature. After blocking, the sections were incubated in the following primary antibodies: mouse monoclonal TNF-α (1: 400, Abcam), rabbit polyclonal IBA1 (Labome, Wako Chemicals USA), rabbit polyclonal GFAP (Dako), chicken polyclonal NeuN (1:500, Saphire bioscience), rabbit polyclonal c-Fos (1:500, Synaptic systems) and their respective secondary antibodies: donkey anti-mouse 647 (1:500, Thermofisher), donkey anti-rabbit 568 (1:500, Invitrogen), goat anti-mouse 488 (1:500, Thermofisher), goat anti-chicken 647 (1:500, Thermofisher). at 4°C overnight. Specifically, 4 sections per brain were stained with IBA1 and GFAP, and another 4 per brain stained for c-Fos and NeuN. Subsequently sections were rinsed with PBS and counterstained with DAPI (Invitrogen) for 10 minutes at room temperature. Finally, the sections were mounted onto SuperFrost slides (Thermofisher Scientific, SuperFrost plus) and coverslipped (Menzel-Glasser, #1) with 50% glycerol mounting medium (Sigma Aldrich).

### Microscopy

Images were captured using TiE2 inverted microscope under 20x and 40x air objective. A z-stack image covering at least 10 µm thickness of the tissue was captured, at 0.6µm/0.9µm stack interval. All images were taken from the dorsal hippocampus which spans from Bregma -1.34 to -2.30 mm based on Paxinos atlas for mouse brain^46^. At least 7 images per brain (left and right hemispheres combined) were obtained for analysis while keeping the laser intensity consistent throughout the image acquisition process. The captured images were then analysed using FIJI (Fiji Is Just ImageJ). For cell counts and morphological analyses, the threshold was adjusted to make sure the cell bodies were all included, (count particles were set at 16 to infinity and this threshold was kept consistent for all sections in CA1. For dentate gyrus it was altered for each image due to inconsistent fluorescent interference from the cell layer). This was followed by intensity measurement, which is represented as Mean grey value (MGV) and background intensity subtracted from the reported MGV. Threshold was not adjusted for intensity measurements. Co-localization was measured for c-Fos and NeuN only, by adjusting the colour threshold to select all the yellow signal (which is the colocalization of red [NeuN] and green[c-Fos]), and the percentage area of the selected signal was measured and averaged for each brain. Thresholds were kept consistent for all sections within the CA1 but were adjusted per image for dentate gyrus (due to fluorescent interference from dense cell layers)”.

## QUANTIFICATION AND STATISTICAL ANALYSIS

The data files and full details of the statistical analyses (i.e. all inputs, outputs, and explanations of outputs) for these experiments can be accessed at the following DOI 10.17605/OSF.IO/6HU7P. The statistical software PSY was used to carry out these analyses. Psy can be downloaded for free onto window-based computers at the following link: https://www.unsw.edu.au/science/our-schools/psychology/our-research/research-tools/psy-statistical-program.

### Cell culture data, Figure 1

For consistency, raw intensity (mean grey value minus background) scores and counts were converted to fold change scores due to the presence of a single control mean (this was not the case for the same measures when quantifying the immunohistochemical data in Figures 4 and 5 due to the presence of separate means for female and male Shams, which if used for fold change calculations, causes a change in the statistical outcomes). Due to the presence of a single control mean, statistical outcomes are identical for cell culture comparisons regardless of whether raw or fold change data was used. All cell culture data was first tested for normality using the Shapiro-Wilks test. If normality was not violated, student t-tests were conducted, if it was violated, Mann-Whitney t-tests were conducted. For all analyses, α = 0.05.

### Behavioural data, Figures 2-3

Lever press and magazine entry data were collected automatically by Med-PC (version 5) and uploaded directly to Microsoft Excel using Med- PC to Excel software. Lever press acquisition and extinction data were analysed using repeated measures (Group × linear) ANOVA controlling the per-family error rate at α = 0.05. For a more fine-grained analysis of test data, we used planned, complex orthogonal contrasts controlling the per-contrast error rate at α = 0.05 according to the procedure described by Hays^98^. If interactions were detected, follow-up simple effects analyses were calculated to determine the source of the interaction. Acquisition data were expressed as mean ± standard error of the mean (SEM) and averaged across counterbalanced conditions. Test data was expressed as each individual data point, overlaying over a histogram representing the mean. Values of p < 0.05 were considered statistically significant.

### Immunohistochemical data, Figures 4-5

Raw intensity (mean grey value minus background) and percentage localisation scores were not converted to fold change scores due to the presence of two separate means for female and male Shams. With such a design, fold change using either a single ‘grand’ mean, or two separate means, alters the nature of statistical analysis. Therefore, raw scores were used. Data were analysed using planned, orthogonal contrasts, controlling the per-contrast error rate at α = 0.05 according to the procedure described by Hays^98^. If interactions were detected, follow-up simple effects analyses were calculated to determine the source of the interaction. Data was expressed as each individual data point, overlaying a histogram representing the mean. Values of p < 0.05 were considered statistically significant.

---

**Supplemental Figure 1.**
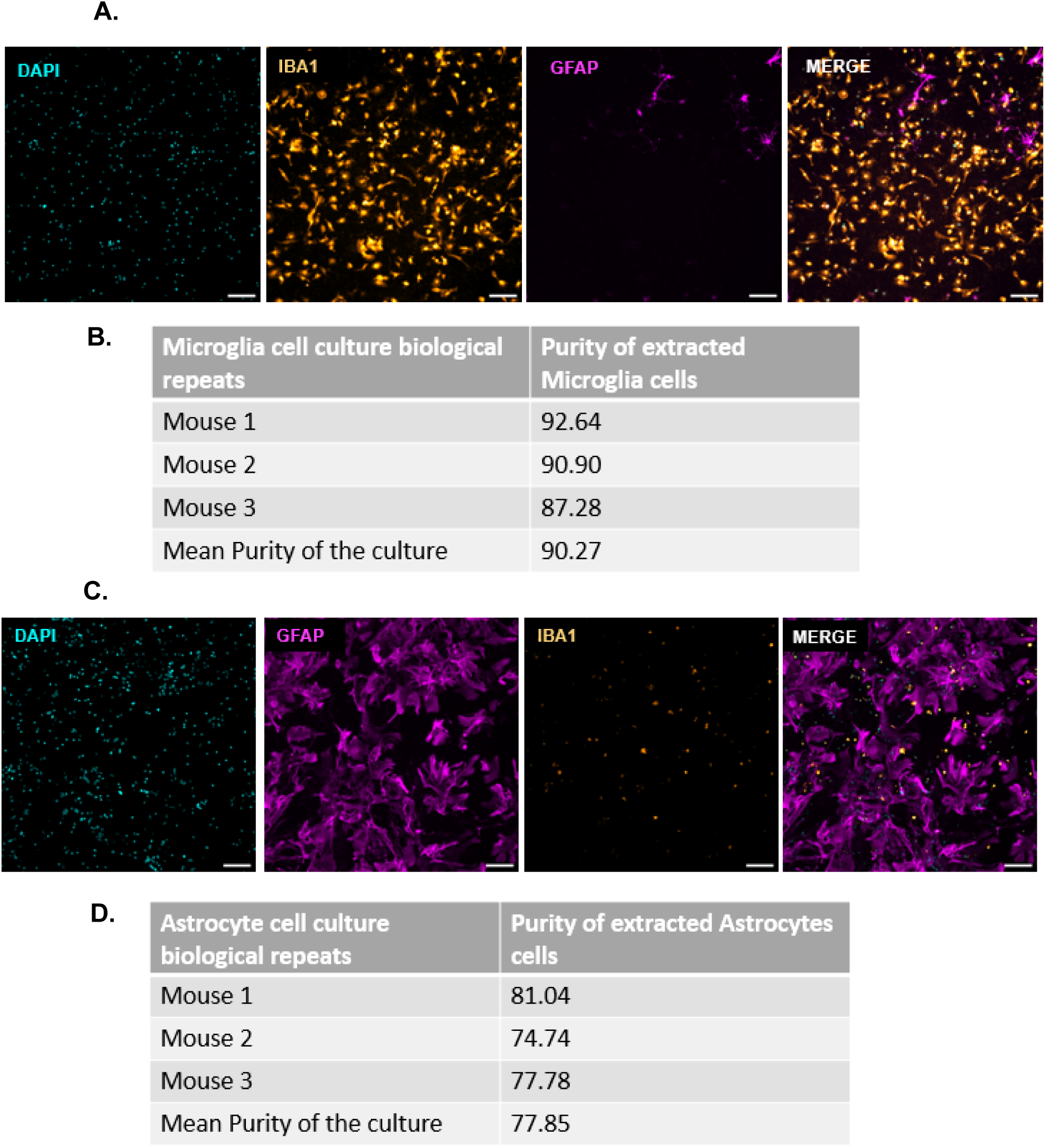
Purity of microglial and astrocytic monocultures. Relates to Figure 1. (A) representative images the microglial monoculture stained with 4ʹ,6-diamidino-2- phenylindole (DAPI)-nucleus, ionised binding calcium adaptor molecule 1 (IBA1)-microglia, glial fibrillary protein (GFAP)-astrocytes, and IBA1/GFAP merged, (B) the calculated purity of the Microglia cell culture from three mouse using Colocalization analysis, (C) representative images of the astrocyte monoculture stained with DAPI, GFAP, IBA1, and GFAP/IBA1 merged, (D) the calculated purity of the astrocytes cell culture from three mouse using colocalization analysis, scale bar = 100µm.

**Supplemental Figure 2:**
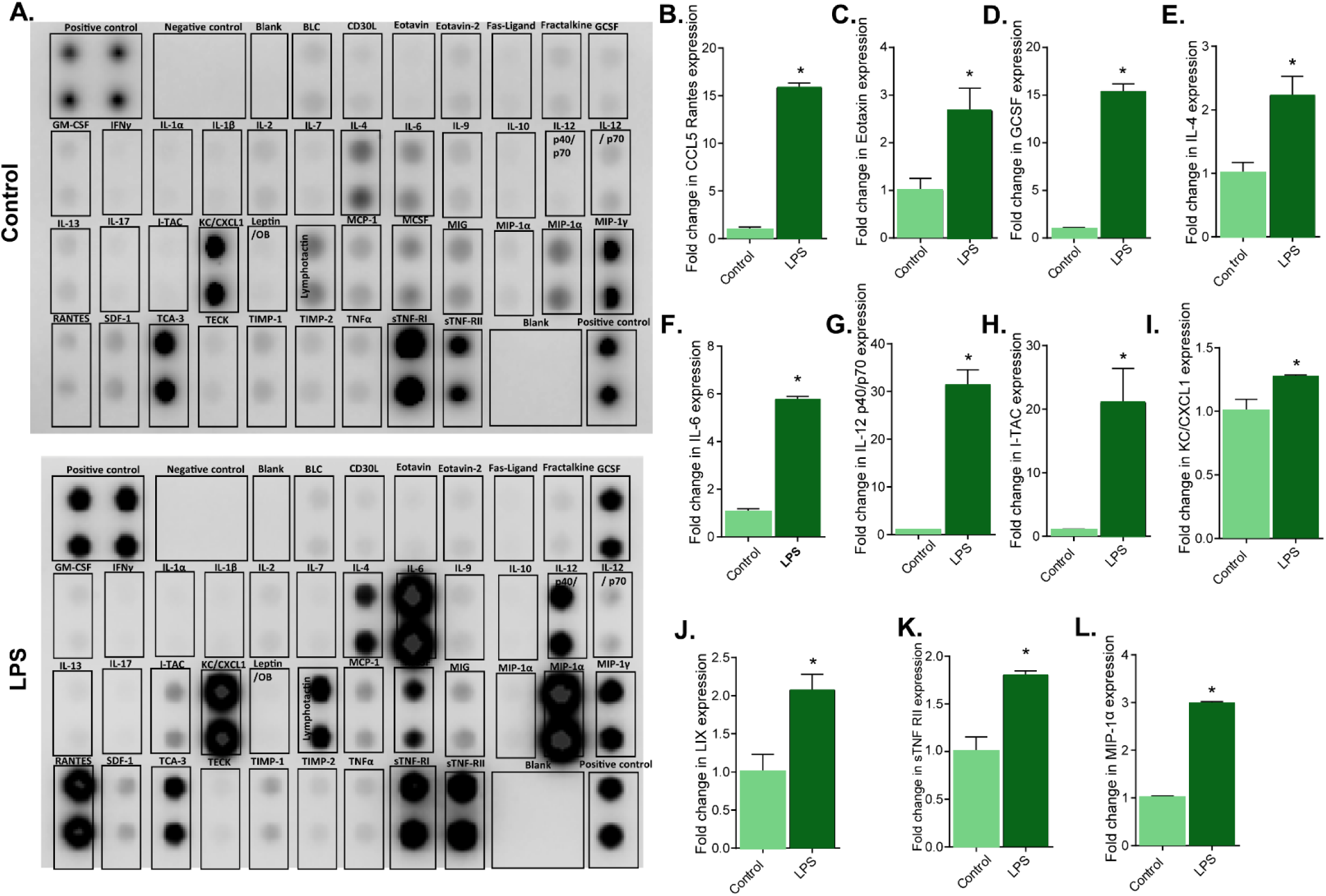
Inflammatory markers from triculture. Supernatants were collected from neuron microglia and astrocyte tricultures and incubated with a mouse inflammation antibody array membrane consisting of 40 mouse inflammatory cytokines. (A) Dot blots, and graphs summarising B) Fold-change of CCL5 rantes, (C) Exotaxin, (D) GCSG, (E) IL-4, (F) IL-6, (G) IL-12 p40/p70, (H) I-TAC, (I) KC/CXCL1, (J) LIX, (K) sTNF-RII, (L) MIP-1α protein expression from tricultures

**Supplemental Figure 3:**
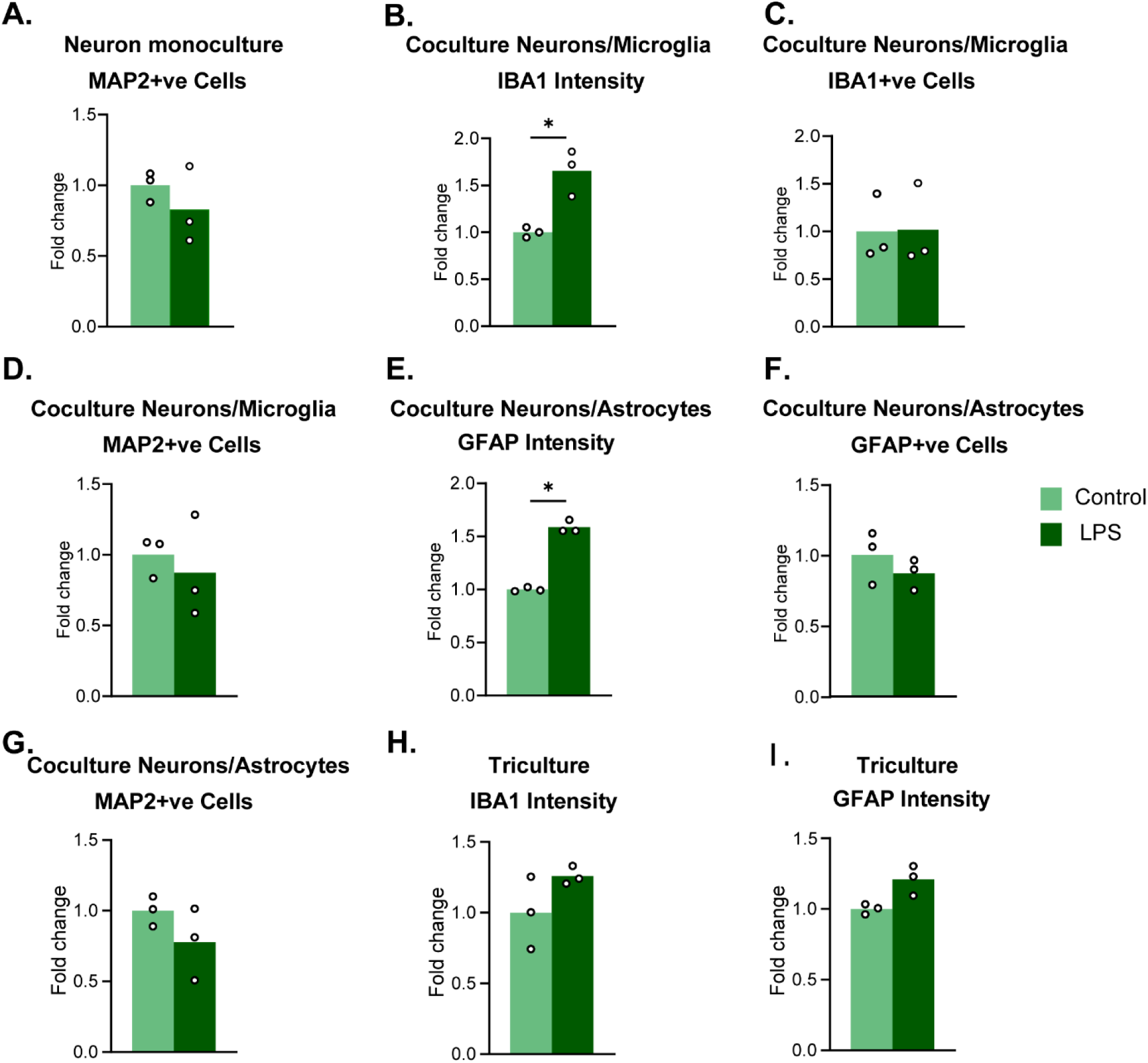
Supplemental cell culture. Relates to Figure 1. (A) Fold change of number of microtubule associated protein 2 (MAP2)+ve cells from neuron monoculture, (B) Fold change of mean grey value measurements for ionised calcium binding molecule 1 (IBA1) in neuron/microglia coculture, (C) Fold change of number of IBA1+vs cells from neuron/microglia coculture, (D) Fold change of number of MAP2+ve cells from neuron/microglia coculture, (E) Fold change of mean grey value measurements for glial acidic fibrillary protein (GFAP) in neuron/astrocyte coculture, (F) Fold change of number of GFAP+ve cells from neuron/astrocyte coculture, (G) Fold change of number of MAP2+ve cells from neuron/astrocyte coculture, (H) Fold change in mean grey value measurements for IBA1 in neuron/microglia/astrocyte triculture, (I) Fold change in mean grey value measurements for GFAP in neuron/microglia/astrocyte triculture. * p < .05

**Supplemental Figure 4:**
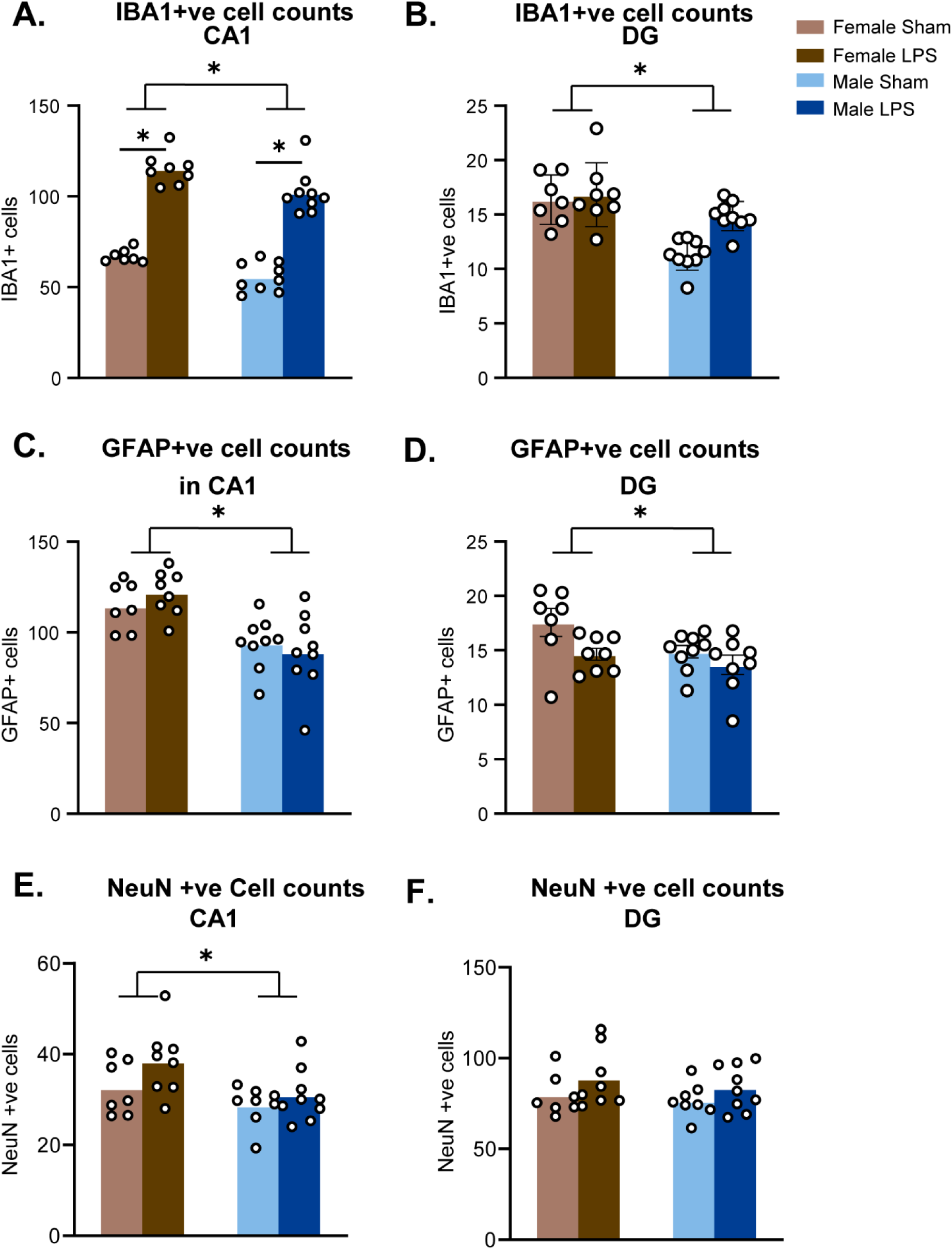
Cell counts from immunohistochemistry. Relates to Figures 4 and 5. (A) number of ionised calcium binding molecule 1 (IBA1)+ve cells in CA1 and, (B) DG regions of hippocampus, (C) number of glial fibrillary acidic protein (GFAP)+ve cells in CA1 and, (D) DG regions of hippocampus, (E) number of NeuN+ve cells in CA1 and, (F) DG regions of hippocampus. * p<.05

